# P granules protect RNA interference genes from silencing by piRNAs

**DOI:** 10.1101/707562

**Authors:** John Paul T. Ouyang, Andrew Folkmann, Lauren Bernard, Chih-Yung Lee, Uri Seroussi, Amanda G. Charlesworth, Julie M. Claycomb, Geraldine Seydoux

## Abstract

P granules are perinuclear condensates in *C. elegans* germ cells proposed to serve as hubs for self/non-self RNA discrimination by Argonautes. We report that a mutant (*meg-3 meg-4*) that does not assemble P granules in primordial germ cells loses competence for RNA-interference over several generations and accumulates silencing small RNAs against hundreds of endogenous genes, including the RNA-interference genes *rde-11* and *sid-1.* In wild-type, *rde-11* and *sid-1* transcripts are heavily targeted by piRNAs, accumulate in P granules, but maintain expression. In the primordial germ cells of *meg-3 meg-4* mutants, *rde-11* and *sid-1* transcripts disperse in the cytoplasm with the small RNA biogenesis machinery, become hyper-targeted by secondary sRNAs, and are eventually silenced. Silencing requires the PIWI-class Argonaute PRG-1 and the nuclear Argonaute HRDE-1 that maintains trans-generational silencing of piRNA targets. These observations support a “safe harbor” model for P granules in protecting germline transcripts from piRNA-initiated silencing.

## Introduction

In the germ cells of animals, dense RNA-protein condensates accumulate on the cytoplasmic face of the nuclear envelope. These condensates, collectively referred to as nuage, contain components of the small RNA (sRNA) machinery that scan germline transcripts for foreign sequences. For example, in *Drosophila*, components of the piRNA machinery in nuage amplify small RNAs that target transcripts from transposable elements for destruction (Huang et al., 2017). In *C. elegans*, the PIWI-class Argonaute PRG-1 associates with ∼15,000 piRNAs encoded in the genome that scan most, if not all, germline mRNAs (Zhang et al., 2018; Shen et al., 2018). PRG-1 accumulates in nuage condensates called P granules that overlay nuclear pores (Batista et al., 2008; Wang and Reinke, 2008). Targeting by PRG- 1/piRNA complexes recruits RNA-dependent RNA polymerases that synthesize 22 nucleotide RNAs (22G- RNAs) complementary to the targeted transcript (Lee et al., 2012; Shen et al., 2018). Synthesis of 22G- RNAs requires proteins in two other nuage condensates: Z granules (ZNFX-1) and mutator foci (MUT-16) that form adjacent to P granules (Ishidate et al., 2018; Wan et al., 2018; Phillips et al., 2012; Zhang et al., 2012). 22G-RNAs in turn are bound by other Argonautes that silence gene expression, including HRDE-1, a nuclear Argonaute that generates a heritable chromatin mark that silences targeted loci for several generations (Buckley et al., 2012). Silencing by exogenous triggers, such as dsRNAs introduced by injection or feeding (exogenous RNAi), also requires 22G-RNA synthesis (Pak and Fire, 2007; Sijen et al., 2007) and HRDE-1 activity, which propagates the RNAi-induced silenced state over generations (Buckley et al., 2012).

The observation that PRG-1/piRNA complexes engage most germline transcripts suggests the existence of mechanisms that restrain PRG-1/HRDE-1 silencing activity (Zhang et al., 2018; Shen et al., 2018). One mechanism involves protection by CSR-1, an opposing Argonaute also present in P granules. CSR-1 binds to abundant 22G-RNAs that target many germline-expressed mRNAs (Seth et al., 2013; Wedeles et al., 2013). CSR-1 opposes the engagement of PRG-1/piRNA complexes (Shen et al., 2018) and is thought to license genes for germline expression (Wedeles et al., 2013; Seth et al., 2013; Cecere et al., 2014; Shen et al., 2018), although some genes are also modestly silenced by CSR-1 (Gerson-Gurwitz et al., 2016). The mechanisms that determine the balance of licensing and silencing 22G-RNAs for each germline-expressed locus are not understood. Inheritance of piRNAs and 22G-RNAs from previous generations is likely to play a role: progeny that inherit neither piRNAs nor 22G-RNAs from their parents and that are competent to synthesize their own 22G-RNAs silence germline genes and become sterile (Phillips et al., 2015; de Albuquerque 2015). P granules could mediate the inheritance of maternal piRNAs and/or 22G-RNAs since P granules contain Argonaute proteins and are maternally inherited (Fig. 1). Segregation of Argonautes and proteins required for 22G-RNA production into distinct nuage compartments (P granules versus Z granules and mutator foci) could also play a role in sorting 22G-RNAs or limiting their production (Wan et al., 2018). A direct test of these hypotheses, however, has been difficult to obtain as complete loss of P granules causes sterility.

**Fig. 1:**
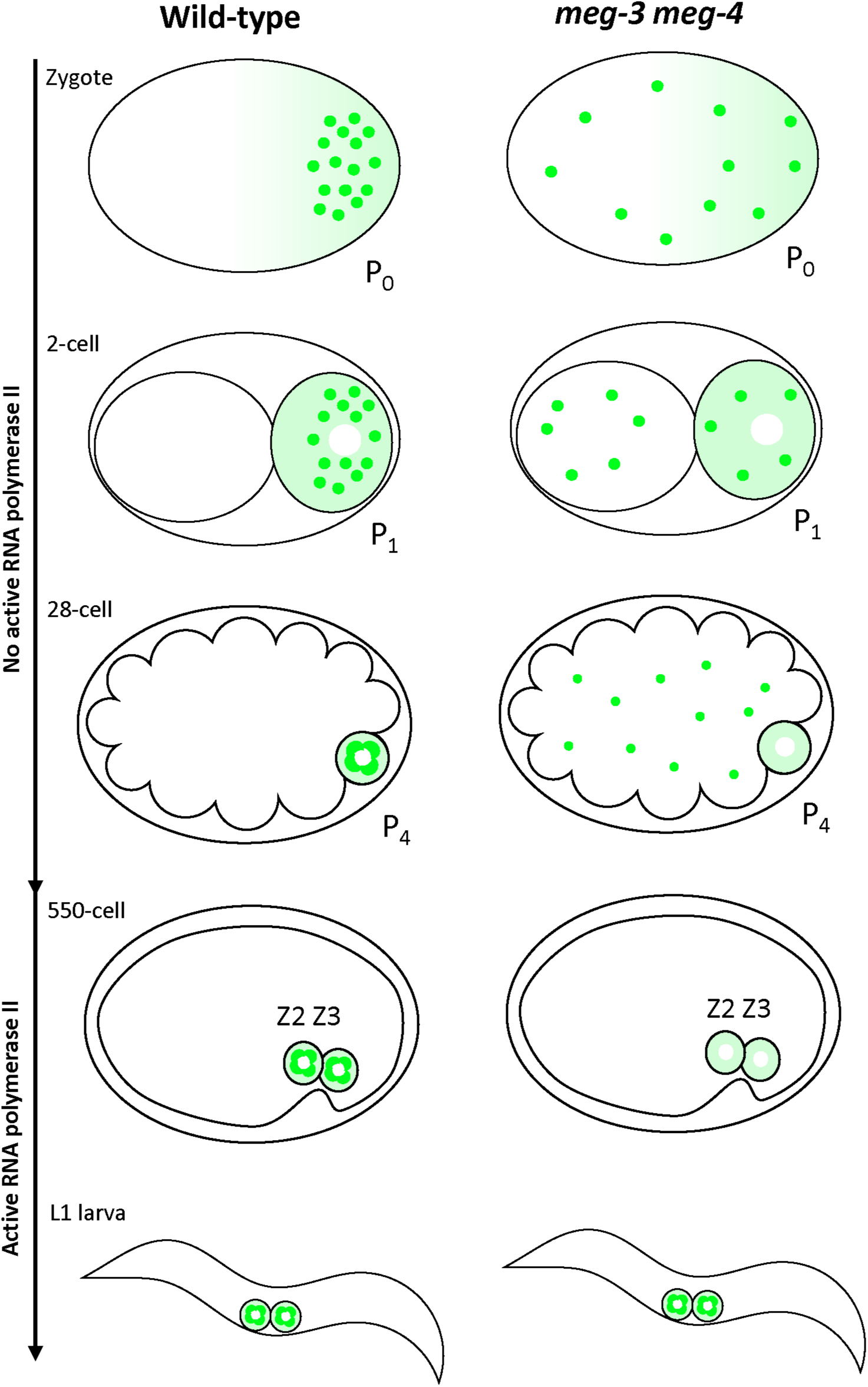
Segregation of P granules in wild-type and *meg-3 meg-4* embryos. Schematics of *C. elegans* embryos at successive stages of development from the 1-cell zygote to the first larval stage post hatching. RNA polymerase II activity is repressed in the P lineage until gastrulation when P_4_ divides to generate Z2 and Z3. In wild-type, P granules (green dots) are segregated preferentially with the germ plasm (lighter green color) to the P lineage that gives rise the primordial germ cells Z2 and Z3. In *meg-3 meg-4* mutants, P granules are partitioned to all cells and are eventually dissolved/turned over. Germ plasm, however, segregates normally in *meg-3 meg-4* mutants. Despite lacking maternal P granules, *meg-3 meg-4* mutants assemble perinuclear P granules *de novo* during late embryogenesis and into the first larval stage (Wang et al., 2014).

We previously identified a mutant that affects P granule coalescence only during embryogenesis (Wang et al., 2014). MEG-3 and MEG-4 are intrinsically-disordered proteins present in the germ plasm, a specialized cytoplasm that is partitioned with the germ lineage during early embryonic cleavages (Wang and Seydoux, 2013). MEG-3 and MEG-4 form gel-like scaffolds that recruit and stimulate the coalescence of P granule proteins in germ plasm to ensure their partitioning to the embryonic germline and the primordial germ cells Z2 and Z3 (Fig. 1; Putnam et al., 2019). In *meg-3 meg-4* embryos, P granules do not coalesce in germ plasm, causing granule components to be partitioned equally to all cells and turned over (Fig. 1; Wang et al., 2014). Despite lacking P granules during embryogenesis, *meg-3 meg-4* assemble P granules *de novo* when the primordial germ cells resume divisions in the first larval stage to generate the ∼ 2000 germ cells that constitute the adult germline. Unlike other P granule mutants, *meg-3 meg-4* mutants are mostly fertile and can be maintained indefinitely (Wang et al., 2014).

In this study, we have examined *meg-3 meg-4* mutants for defects in small RNA (sRNA) homeostasis. We find that *meg-3 meg-4* mutants become progressively deficient in exogenous RNA-mediated interference over several generations and accumulate abnormally high levels of sRNAs that silence endogenous genes. The silenced genes belong to a class of genes that in wild-type are targeted primarily by the silencing Argonautes PRG-1 and HRDE-1, and include *rde-11* and *sid-1*, two genes required for exogenous RNAi. *rde-11* and *sid-1* transcripts are retained in P granules in wild-type, but in *meg-3 meg-4* mutants, the transcripts become dispersed in the cytoplasm with Z granules and mutator foci components. Our findings suggest a role for P granules in protecting certain germline transcripts from run-away, trans-generational silencing initiated by piRNAs and amplified by HRDE-1-associated 22Gs.

## Results

### *meg-3 meg-4* mutants are defective in exogenous RNA-mediated interference

JH3475 is a strain in which both the *meg-3* and *meg-4* open reading frames have been deleted by genome editing (Smith et al., 2016; Paix et al., 2017). This strain (*meg-3 meg-4^#1^*) has been passaged over 100 times. In the course of conducting experiments with *meg-3 meg-4^#1^* worms, we noticed that *meg-3 meg-4^#1^* adults appeared resistant to exogenous RNA-mediated interference. To examine this phenotype systematically, we fed *meg-3 meg-4^#1^* hermaphrodites bacteria expressing double-stranded RNA (dsRNA) against the *pos-1* gene. *pos-1* is a maternally-expressed gene required for embryonic viability (Tabara et al., 1999). As expected, wild-type control hermaphrodites laid on average only 6.5% viable embryos after *pos-1(RNAi)* (Fig. 2A). In contrast, *meg-3 meg-4^#1^* laid on average 76% viable embryos after *pos-1(RNAi)* (Fig. 2A). We obtained similar results by administering the double-stranded RNA by injection, and by targeting two other maternally-expressed genes required for embryogenesis (*mex-5 and mex-6*) (Fig. 2A and S1A). Abnormal RNAi behavior of strains with loss of function mutations in *meg-3* and *meg-4* has also been reported by others (Wan et al., 2018; Lev et al., 2019).

**Fig. 2:**
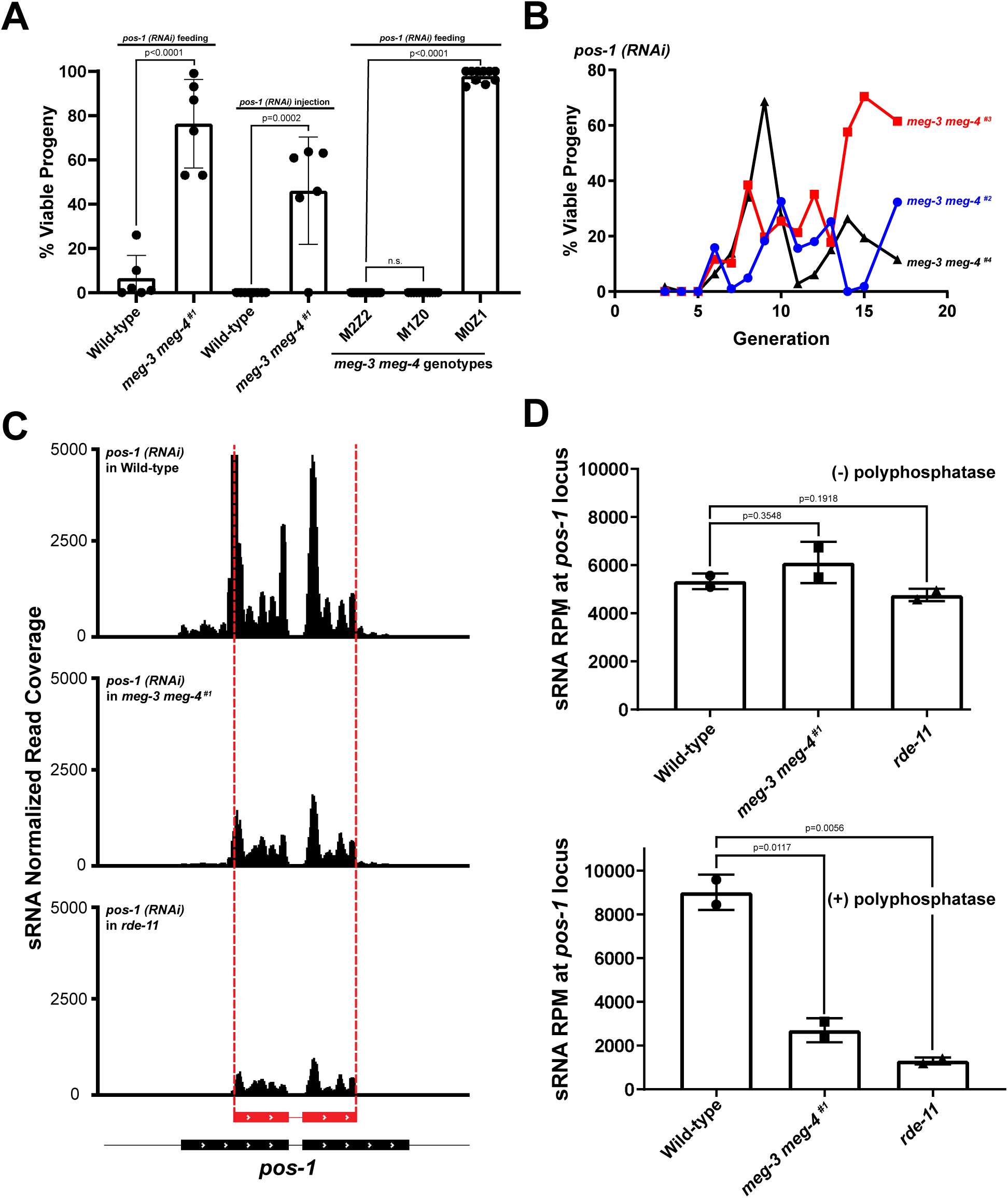
*meg-3 meg-4* mutants lose competency for RNA-interference and are defective in the production of secondary siRNAs. A. Graph showing the percentage of viable embryos laid by hermaphrodites of the indicated genotypes upon treatment with *pos-1* dsRNA. First two bars depict the embryonic viability from populations of ∼20 hermaphrodites fed starting at the L1 stage (each dot represents an experiment performed on a distinct population). On average, roughly 200 embryos were scored per RNAi experiment. The following two bars represent the percent viable progeny of mothers ∼16 hours following injection with 200 ng/uL of *pos-1* dsRNA (each dot represents the progeny of a single injected young adult hermaphrodite that laid more than 15 embryos). The last three bars represent viable progeny from M2Z2, M1Z0, and M0Z1 hermaphrodites fed starting at the L4 stage (each dot represents the progeny of a single hermaphrodite that laid more than 15 embryos). The “M” and “Z” designations refer to the number of wild-type *meg-3 meg-4* alleles present in the mother (M) or hermaphrodite (Z) tested for RNAi. Bar height represents the mean; error bars represent the standard deviation. P-values were calculated using an unpaired t-test. B. Graph showing the percentage of viable embryos among broods (∼12 mothers) laid by newly generated *meg-3 meg-4* hermaphrodites fed with bacteria expressing *pos-1* dsRNA (from L4 stage). Three independently derived strains are shown. “Generation” refers to the number of generations since the *meg-4* gene was deleted by genome editing in the starting strain carrying only a *meg-3* deletion. See Fig. S1C for RNAi sensitivity of three sibling strains carrying only the original *meg-3* deletion. See Fig. S1D for CRISPR breeding scheme. C. Genome browser view of sRNA reads mapping to the *pos-1* locus in adult hermaphrodites of indicated genotypes fed with bacteria expressing a dsRNA trigger (red in figure) against a central region of the *pos-1* locus. D. Graphs showing the abundance of sRNA reads mapping to the *pos-1* locus in adult hermaphrodites of the indicated genotypes fed *pos-1* RNAi. The upper panel shows primary sRNAs (directly derived from the ingested trigger), the bottom graph shows all sRNAs (both primary and secondary) from phosphatase treated library samples. Bar height represents the mean; error bars represent the standard deviation; p-values were calculated using an unpaired t-test.

*meg-3* and *meg-4* are required maternally for the formation of P granules in embryos (Wang et al., 2014). To determine whether *meg-3* and *meg-4* were also required maternally for RNAi competence, we tested *meg-3 meg-4* homozygous hermaphrodites derived from heterozygous *meg-3 meg-4^#1^/++* mothers (M1Z0) and *meg-3 meg-4^#1^/++* heterozygous hermaphrodites derived from homozygous mutant mothers (M0Z1) (see Fig. S1B for crosses). We found that M1Z0 hermaphrodites had normal sensitivity to RNAi, whereas M0Z1 hermaphrodites were defective, consistent with a maternal requirement for *meg-3 meg-4* (Fig. 2A). To test this further, using genome editing (Paix et al., 2017), we regenerated the *meg-4* deletion in a line carrying the *meg-3* deletion to generate three new *meg-3 meg-4* lines (*meg-3 meg-4 ^#2^*, *meg-3 meg-4 ^#3^*, and *meg-3 meg-4 ^#4^*). Strikingly, we found that the newly generated *meg-3 meg-4* lines remained competent for RNAi for at least five generations before beginning to exhibit resistance. After generation six, the degree of RNAi resistance varied from generation to generation and between strains (Fig. 2B). In contrast, three sibling strains that only contained the *meg-3* deletion remained sensitive to RNAi throughout the course of the experiment (Fig. S1C). We conclude that *meg-3 meg-4* mutants exhibit a defect in RNAi that is acquired progressively over several generations.

### *meg-3 meg-4* mutants exhibit reduced accumulation of secondary siRNAs triggered by *pos-1(RNAi)*

Silencing of gene activity after ingestion of a long double-stranded RNA trigger requires production of primary sRNAs derived from the trigger, and synthesis of secondary sRNAs templated from the targeted RNA (Yigit et al., 2006; Pak and Fire, 2007; Sijen et al., 2007). To determine which step is affected in *meg-3 meg-4* mutants, we sequenced sRNAs from wild-type and *meg-3 meg-4^#1^* adult hermaphrodites fed bacteria expressing *pos-1* dsRNA. As an additional control, we also sequenced sRNAs from *rde-11* hermaphrodites fed *pos-1* RNAi bacteria. *rde-11* mutants generate primary sRNAs but fail to generate secondary sRNAs and are defective in exogenous RNAi (Yang et al., 2012; Zhang et al., 2012). Primary and secondary sRNAs can be differentiated by the presence of a 5’ monophosphate on primary sRNAs and a 5’ triphosphate on secondary sRNAs (Pak and Fire, 2007; Sijen et al., 2007). Therefore, for each genotype, we prepared two types of libraries: one where the RNA was left untreated to preferentially clone primary siRNAs and one where the RNA was treated with a 5’ polyphosphatase to allow the cloning of both primary and secondary sRNAs. As expected, we found that wild-type hermaphrodites accumulate many sRNAs at the *pos-1* locus that target sequences both within and outside the trigger (Fig. 2C). *rde-11* mutants in contrast accumulate fewer sRNAs at the *pos-1* locus and all of these target sequences within the trigger region, consistent with normal production of primary sRNAs and defective production of secondary sRNAs as reported previously (Fig. 2C and Zhang et al., 2012). Similar to *rde-11*, *meg-3 meg-4* mutants accumulated fewer sRNAs at the *pos-1* locus, and these sRNAs mapped primarily to the trigger (Fig. 2C). Quantification of primary sRNAs at the *pos-1* locus revealed similar levels of primary sRNAs in all genotypes (no treatment samples), and reduced overall levels of sRNAs in *rde-11* and *meg-3 meg-4* compared to wild-type (5’ polyphosphatase-treated samples) (Fig. 2D). We conclude that, like *rde-11* mutants, *meg-3 meg-4* mutants are defective in the production of secondary sRNAs generated in response to an exogenous RNA trigger.

### *meg-3 meg-4* mutants have elevated numbers of sRNAs against *rde-11* and five other genes implicated in small RNA pathways

MEG-3 and MEG-4 proteins are expressed primarily in embryos (Fig. S2A), and so are unlikely to have a direct role in the production of secondary sRNAs in larval and adult hermaphrodites. The generational delay in the appearance of the RNAi defective phenotype also suggests an indirect effect. To understand the origin of the RNAi defect in *meg-3 meg-4* mutants, we sequenced sRNAs in mixed populations of *meg-3 meg-4^#1^*, *meg-3 meg-4 ^#2^*, *meg-3 meg-4 ^#3^*, and *meg-3 meg-4 ^#4^* under normal feeding conditions (no exogenous RNAi). We considered three classes of sRNAs: piRNAs and microRNAs, which are genomically encoded, and sRNAs that are antisense to coding genes. The latter can be sub-divided further based on published lists of sRNAs immunoprecipitated with specific Argonautes (Methods). We detected all major classes of sRNAs in *meg-3 meg-4* mutants, including piRNAs, microRNAs and sRNAs mapping to loci targeted by the Argonautes WAGO-1, WAGO-4, HRDE-1, and CSR-1 (Fig. S2B; Gu et al., 2009; Xu et al., 2018, Buckley et al., 2012; Claycomb et al., 2009). All classes accumulated at levels similar to wild-type, with the exception of microRNAs which appeared slightly elevated in *meg-3 meg-4* mutants (Fig. S2B). We also compared the sRNA length distribution and 5’ nucleotide preference in wild-type and *meg-3 meg-4^#1^* sRNA libraries and found no overt differences (Fig. S2C-D).

We compared the frequency of sRNA reads at every annotated locus in the genome in *meg-3 meg-4* mutants versus wild-type. Surprisingly, we identified hundreds of loci with misregulated sRNAs (Fig. 3A and Fig. S2E-G). Combining data for all four strains, we identified 303 and 316 loci that were targeted by more or fewer sRNAs, respectively, in all four strains compared to wild-type (Tables S1-S2). Interestingly, nearly 50% of those loci have been reported to be targeted by sRNAs associated with HRDE-1 in wild-type hermaphrodites (Fig. 3B). HRDE-1-associated sRNAs target 1,208 loci in wild-type, and 25% (306) of those loci exhibit mis-regulated sRNAs in *meg-3 meg-4* mutants (Fig. 3C). In contrast, CSR-1-associated sRNAs target over 4,000 transcripts, but only 1.2% (50) of these exhibited misregulated sRNAs in *meg-3 meg-4* mutants (Fig. 3C). We conclude that *meg-3 meg-4* mutants misregulate sRNA at many loci that are primarily targeted by the silencing Argonaute HRDE-1.

**Fig. 3:**
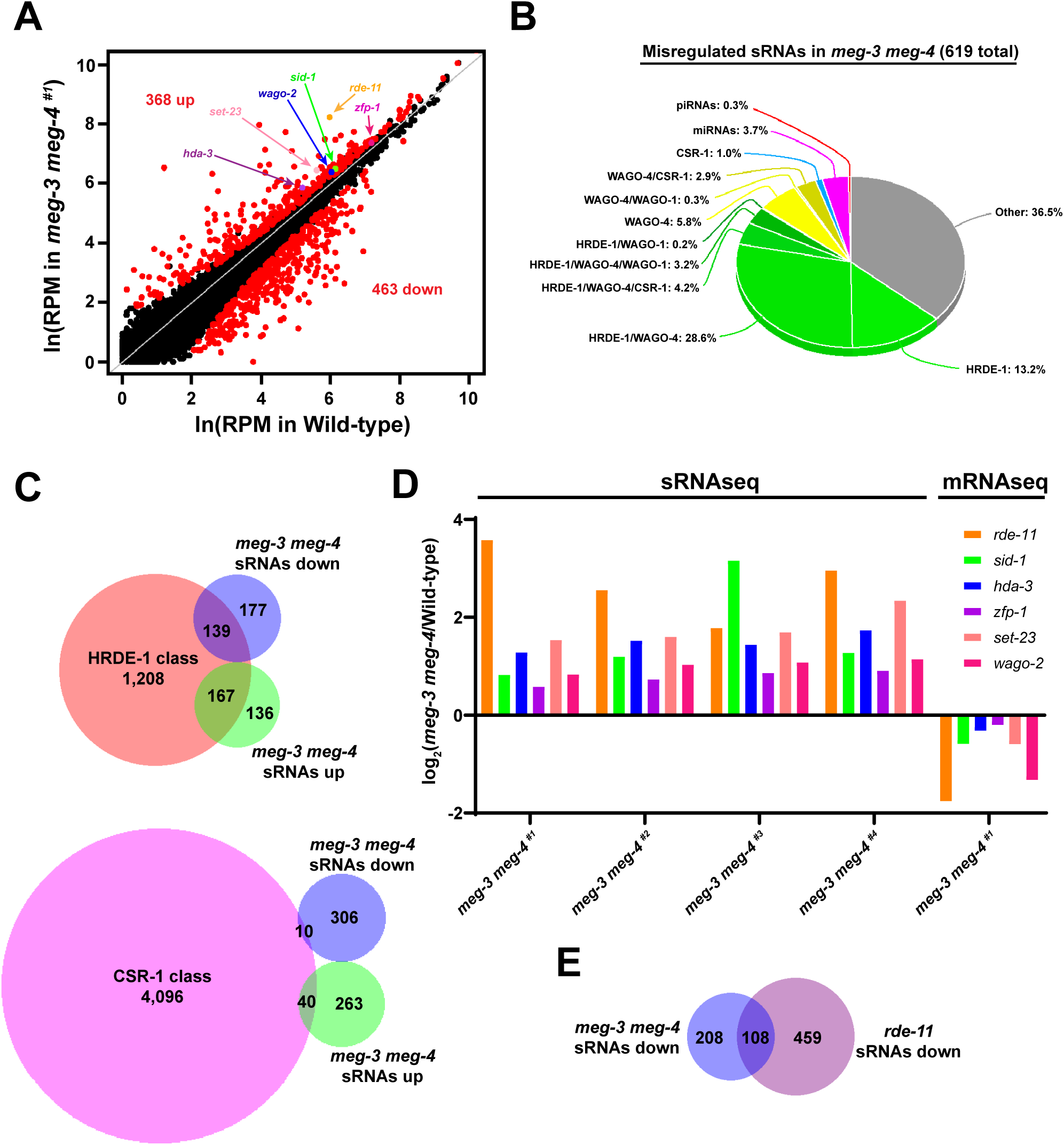
*meg-3 meg-4* mutants misregulate sRNAs that target hundreds of loci. A. Scatter plot comparing sRNA abundance in wild-type (X-axis) and *meg-3 meg-4^#1^*(Y-axis) hermaphrodites. Each dot represents an annotated locus in the *C. elegans* genome. Red dots represent loci with significantly upregulated or downregulated sRNAs comparing two biological replicates each for wild-type and *meg-3 meg-4^#1^*. B. Pie chart showing the 619 genes with misregulated sRNAs in *meg-3 meg-4* strains categorized according to the type of sRNAs that target these genes in wild-type. Note that 49.4% of these sRNAs are classified as HRDE-1-associated (Buckley et al., 2012). C. Venn diagrams showing the overlap between loci with upregulated or downregulated sRNAs in *meg-3 meg-4* mutants and loci targeted by sRNAs that co-immunoprecipitate with HRDE-1 and CSR-1 (Buckley et al., 2012; Claycomb et al., 2009). D. Bar graph showing the average log2 fold difference in sRNA abundance for the indicated loci in the four *meg-3 meg-4* strains compared to wild-type. The log2 fold change represents the average of two biological replicates for each genotype. Last grouping shows the mRNA abundance for each gene in the *meg-3 meg-4^#1^* adults as determined by RNAseq from two biological replicates. E. Venn diagrams showing the overlap between loci with downregulated sRNAs in *meg-3 meg-4* mutants and loci with downregulated sRNAs in *rde-11* mutants.

We reasoned that upregulation of silencing sRNAs against loci required for RNAi could explain the RNAi defective phenotype of *meg-3 meg-4* mutants. To investigate this possibility, we cross-referenced the 303 genes with upregulated sRNAs with a list of 332 genes implicated in small RNA pathways compiled from the “gene silencing by RNA” Gene Ontology classification of WormBase WS270, Kim et al., 2005, and Tabach et al., 2013 (Table S3). This analysis identified 6 genes: *rde-11*, *sid-1*, *hda-3*, *zfp-1*, *set-23* and *wago-2. rde-11* codes for a RING finger domain protein required for exogenous RNAi as described above (Yang et al., 2012; Zhang et al., 2012). *sid-1* codes for a dsRNA transporter required for exogenous RNAi by feeding (Winston et al., 2002; Feinberg and Hunter 2003; Minkina and Hunter, 2017). *hda-3* and *zfp-1* are chromatin factors identified in a screen for genes required for exogenous RNAi (Kim et al., 2005). *set-23* is a predicted histone methyltransferase identified in a screen for genes that co-evolved with known RNAi factors (Tabach et al., 2013). *wago-2* is a member of the 27 Argonautes present in the *C. elegans* genome (Yigit et al., 2006) and a predicted pseudogene (WormBase WS270). sRNAs against the six genes were elevated in all four strains, but the extent of upregulation varied from strain to strain and gene to gene, with *rde-11* and *sid-1* showing the highest increase in three and one of the four strains, respectively (Fig. 3D). We reasoned that elevated sRNAs might result in downregulation of the corresponding mRNA transcript. For those analyses, we used *meg-3 meg-4^#1^*, the oldest *meg-3 meg-4* strain with a strong RNAi-resistant phenotype. We found that expression of the six genes appeared reduced in *meg-3 meg-4^#1^*compared to wild-type as determined by RNAseq (Fig. 3D – the difference for *zfp-1* did not score as statistically significant). The RNAseq data, however, must be interpreted cautiously since RNAseq was performed on populations of adult worms, which in the case of *meg-3 meg-4^#1^* include ∼ 30% worms lacking a germline (Wang et al., 2014), but see below for a more direct measurement of *rde-11* transcript levels. Together, these observations suggest that the RNAi defect of *meg-3 meg-4* mutants is caused by increased targeting by sRNAs (and likely lower mRNA expression) of 4 genes with a demonstrated requirement in exogenous RNAi (*rde-11*, *sid-1*, *hda-3,* and *zfp-1)* and two genes (*set-23* and *wago-2*) with potential roles in sRNA pathways.

We also cross-referenced the sRNAs downregulated in *meg-3 meg-4* with sRNA pathway genes and identified only one gene (*haf-4*). Expression of this gene did not change significantly in *meg-3 meg-4^#1^*. We noticed, however, that 34% of loci with downregulated sRNAs in *meg-3 meg-4* mutants also exhibited downregulated sRNAs in *rde-11* mutants (Fig. 3E). This observation suggests that downregulation of some sRNAs in *meg-3 meg-4* mutants may be an indirect consequence of reduced *rde-11* activity.

### The nuclear Argonaute *hrde-1* is required for upregulation of sRNAs at the *rde-11* and *sid-1* loci in *meg-3 meg-4* mutants

Nearly 50% of the genes with misregulated sRNAs in *meg-3 meg-4* mutants (306 of 619 genes) are targeted by HRDE-1-associated sRNAs in wild-type (Fig. 3B). HRDE-1 is a nuclear Argonaute that recruits the nuclear RNAi machinery to nascent transcripts. Interestingly, we noticed that the distribution of sRNAs mapping to the *rde-11* locus in *meg-3 meg-4* mutants is consistent with silencing by the nuclear RNAi machinery. *rde-11* is transcribed as part of an operon that includes *B0564.2,* a gene immediately 3’ of *rde-11*. Operons are transcribed as single, long transcripts that are broken up into shorter transcripts by trans-splicing in the nucleus before transport to the cytoplasm (Blumenthal and Gleason, 2003). In wild-type, only exons three and four of *rde-11* were targeted by sRNAs, with fewer sRNA mapping to the other exons of *rde-11* or to *B0564.2.* In contrast, in *meg-3 meg-4* mutants, all exons of both genes were heavily targeted by sRNAs (Fig. 4A and Fig. S3A). As observed for *rde-11, B0564.2* mRNA levels were also significantly downregulated in *meg-3 meg-4^#1^* as determined by RNAseq (Fig. S3A). The observation that *rde-11* and *B0564.2* are co-targeted by small RNAs in *meg-3 meg-4* mutants is consistent with targeting by a nuclear Argonaute (Guang et al., 2008).

**Fig. 4:**
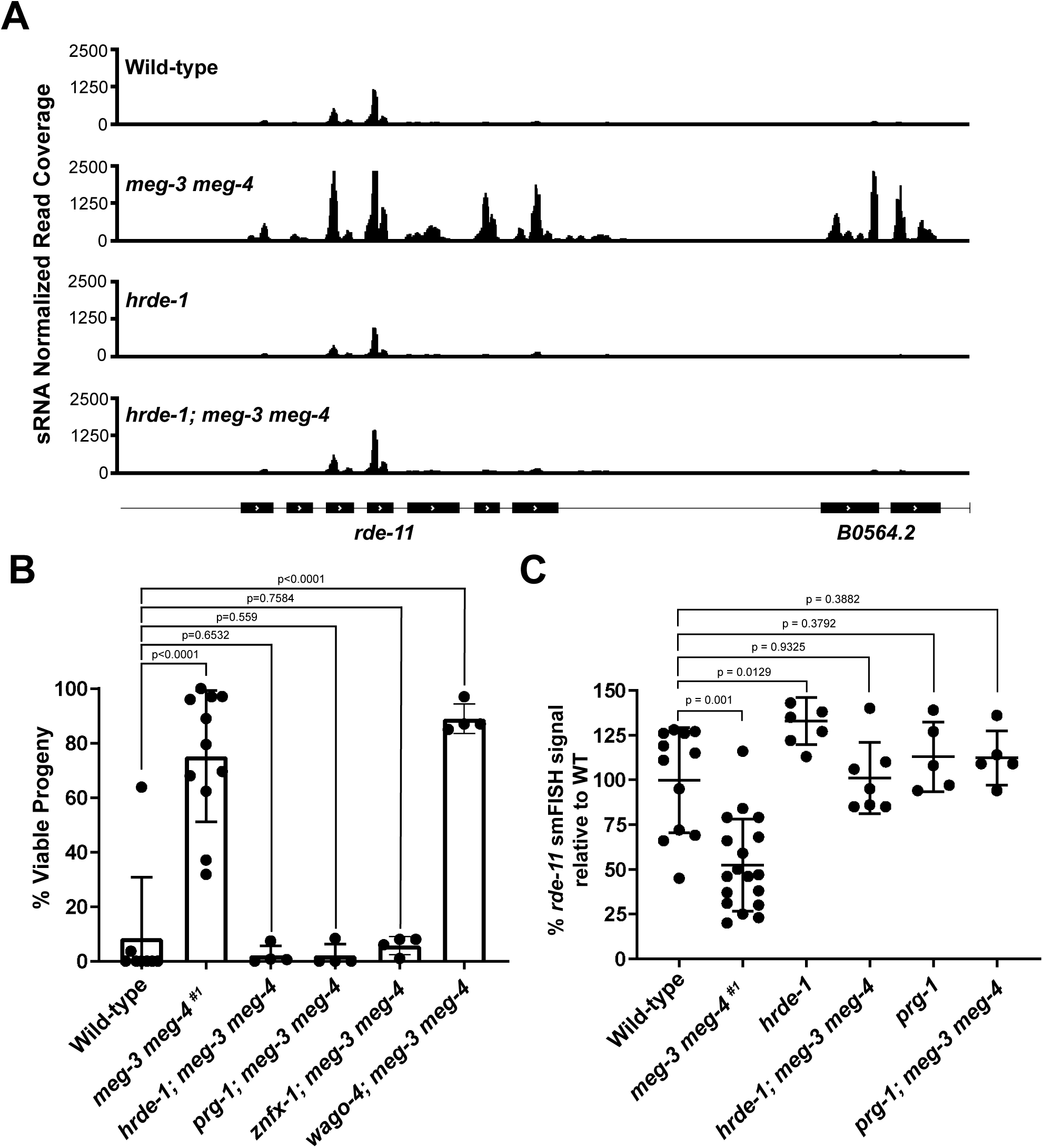
*meg-3 meg-4* phenotypes are suppressed by loss-of-function mutations in *hrde-1* and *prg-1*. A. Browser view of the *rde-11/B0564.2* locus showing normalized sRNA reads in hermaphrodites of the indicated genotypes. B. Graph showing the percentage of viable embryos among broods laid by hermaphrodites of the indicated genotypes and fed bacteria expressing *pos-1* dsRNA from the L1 stage. Each dot represents an independent RNAi experiment performed with a cohort of 15-20 hermaphrodites allowed to lay eggs for 1-2 hours. On average, over 200 embryos were scored per RNAi experiment. Note for *prg-1; meg-3 meg-4*, values were normalized to the levels of embryonic lethality the strain exhibits under non-RNAi conditions. Bar height and error bars represent the mean and standard deviation respectively; p-values were obtained using an unpaired t-test. C. Quantification of smFISH signal normalized to the average wild-type value. Each dot represents a single gonad. Center bar represents the mean and error bars indicate the standard deviation. P values were obtained through an unpaired t-test. See Fig S3G for regions quantified.

We reasoned that if HRDE-1 were required for silencing the *rde-11* operon, a loss of function mutation in *hrde-1* should block sRNA amplification against the *rde-11* and *B0564.2* loci and restore transcripts levels back to wild-type. To test this, we crossed *meg-3 meg-4* hermaphrodites with males carrying a mutation in *hrde-1* to generate the triple mutant *hrde-1; meg-3 meg-4* (see Fig. S3B for crosses). Consistent with our hypothesis, we observed lower levels of sRNAs against the *rde-11* and *B0564. 2* transcripts in *hrde-1; meg-3 meg-4* compared to *meg-3 meg-4* (Fig. 4A). sRNAs against *sid-1,* were also significantly reduced (Fig. S3C), whereas sRNAs against the other sRNA pathway genes (*wago-2*, *hda-3*, *set-23*, and *zfp-1*) did not show changes that reached statistical significance (Fig. S3C). Of the 303 transcripts with upregulated sRNAs in *meg-3 meg-4* mutants, only 39 were partially rescued (lowered) in *hrde-1; meg-3 meg-4* (Table S4). Although this analysis is likely to be complicated by sRNA defects inherent to loss of *hrde-1* activity, we conclude that *hrde-1* is responsible for some, but not all, of the upregulation of sRNAs in *meg-3 meg-4* mutants. Other Argonautes that overlap in function with HRDE-1 may be responsible for the remainder (Shirayama et al., 2012; Gu et al., 2009).

### *rde-11* and *sid-1* are engaged by PRG-1-piRNA complexes and not by CSR-1-sRNA complexes

HRDE-1 has been shown to act downstream of the piRNA Argonaute PRG-1 to perpetuate a sRNA epigenetic memory (Ashe et al., 2012; Shirayama et al., 2012). Using previously published Cross Linking and Selection of Hybrids (CLASH) data (Shen et al., 2018), we assigned a rank to each protein coding gene based on degree of targeting by PRG-1/piRNA complexes. We found that *rde-11* and *sid-1* rank among the top 50 genes in the genome most targeted by PRG-1/piRNAs complexes (average rank among coding genes across two CLASH replicates: #15 for *rde-11*, *#*33 for *sid-1*). 123 unique piRNA sites were identified in the *rde-11* transcript and 75 in the *sid-1* transcript (Shen et al., 2018). Consistent with targeting by piRNAs, sRNAs targeting *rde-11* and *sid-1* were reduced in *prg-1* mutants as compared to wild-type whereas *rde-11* and *sid-1* mRNA levels were increased in *prg-1* mutants (Lee et al., 2012; Shen et al., 2018; McMurchy et al., 2017, Fig. S3D-E). Silencing of endogenous genes by PRG-1 is countered by the Argonaute CSR-1, which licenses germline genes for expression (Wedeles et al., 2013; Seth et al., 2013; Cecere et al., 2014; Shen et al., 2018). Interestingly, a published list of sRNAs that co-immunoprecipitate with CSR-1 did not contain sRNAs against *rde-11* or *sid-1* (Claycomb et al., 2009). In fact, as noted above, more than 90% of loci with misregulated sRNAs in *meg-3 meg-4* mutants do not appear to be targeted by CSR-1-associated sRNAs (Fig. 3B). These observations suggest that misregulated genes in *meg-3 meg-4* mutants may be in a “sensitized” state in wild-type: hyper-targeted by silencing PRG-1/piRNA complexes and hypo-targeted by protective CSR-1/sRNA complexes.

### PRG-1 and HRDE-1 are required for *rde-11* silencing and for the RNAi-defective phenotype of *meg-3 meg-4* mutants

We reasoned that if PRG-1 and HRDE-1 are responsible for the hyper-targeting of loci required for exogenous RNAi in *meg-3 meg-4* mutants, loss of function mutations in *prg-1* and *hrde-1* should restore competence for exogenous RNAi to *meg-3 meg-4* mutants. As predicted, we found that, unlike *meg-3 meg-4* mutants, *hrde-1; meg-3 meg-*4 and *prg-1; meg-3 meg-4* mutants were competent for RNAi (Fig. 4B; see Fig. S3F for cross). In contrast, mutations in a different Argonaute WAGO-4 did not suppress the *meg-3 meg-4* phenotype (Fig. 4B; see Fig. S3F for cross). CSR-1 mutants are sterile and so could not be tested in this assay (Yigit et al., 2006; Claycomb et al., 2009). ZNFX-1 is a conserved helicase required for sRNA amplification (Ishidate et al., 2018; Wan et al., 2018). We found that *znfx-1; meg-3 meg-4* worms were competent for RNAi, suggesting that ZNFX-1, like PRG-1 and HRDE-1, is required for hypertargeting of RNAi loci in *meg-3 meg-4* mutants (Fig 4B; see Fig. S3F for cross).

To examine whether *rde-11* expression is restored in *meg-3 meg-4* mutants that also lack *prg-1* or *hrde-1*, we used single-molecule fluorescent *in situ* hybridization to directly measure *rde-11* transcript levels in adult germlines. We focused on *rde-11* since that locus showed the greatest reduction in mRNA level in a population of *meg-3 meg-4^#1^* adults (Fig. 3B). We found that, as expected, *rde-11* is expressed robustly in wild-type germlines and at much lower levels in *meg-3 meg-4^#1^*germlines (Fig. 4C and Fig. S3G). Remarkably, wild-type levels of *rde-11* transcripts were restored in *hrde-1; meg-3 meg-4* and *prg-1; meg-3 meg-4* germlines (Fig. 4C and Fig. S3G). We conclude that PRG-1 and HRDE-1 are required for silencing of the *rde-11* locus in *meg-3 meg-4* adult germlines.

### P granule proteins, including PRG-1, fail to coalesce into granules in *meg-3 meg-4* embryos

Previous studies using the P granule marker PGL-1 showed that P granules assemble normally post-embryogenesis in *meg-3 meg-4* germlines (Wang et al., 2014). We verified this observation and confirmed that formation of Z granules and mutator foci was also unaffected in adult *meg-3 meg-4* germlines (Fig. S4A-B, Wan et al., 2018). Additionally, PRG-1 and CSR-1 protein levels appeared unchanged in *meg-3 meg-4* adults compared to wild-type as determined by western analyses (Fig. S4C). Silencing of the *rde-11* locus in adult germlines, therefore, is unlikely to be due to gross defects in nuage organization at this stage.

During the oocyte-to-embryo transition, the canonical P granule component PGL-1 relocalizes from the nuclear periphery to cytoplasmic granules that are asymmetrically partitioned to the embryonic germ lineage during the first embryonic cleavages (Strome and Wood, 1982). Whether other nuage components behave similarly has not yet been reported systematically. Using fluorescently-tagged alleles generated by genome editing, we compared the distribution of PRG-1, CSR-1, ZNFX-1, and MUT-16 to that of PGL-1 (Fig. 5A; Methods, Shen et al., 2018, Wan et al., 2018). We found that like PGL-1, PRG-1 and ZNFX-1 localize to granules that segregate preferentially with the germ lineage during early cleavages (also see Wan et al., 2018). CSR-1 exhibited a similar pattern, except that CSR-1 granules did not appear as strongly asymmetrically segregated (Fig 5A). Around the 28-cell stage, PGL-1 becomes concentrated in autophagic bodies in somatic cells and is turned over (Zhang et al., 2009). We observed a similar pattern of turnover for PRG-1, CSR-1 and ZNFX-1 in somatic lineages. By comma-stage, PGL-1, PRG-1, CSR-1 and ZNFX-1 could only be detected in the primordial germ cells Z2 and Z3 (Fig. 5A).

**Fig. 5:**
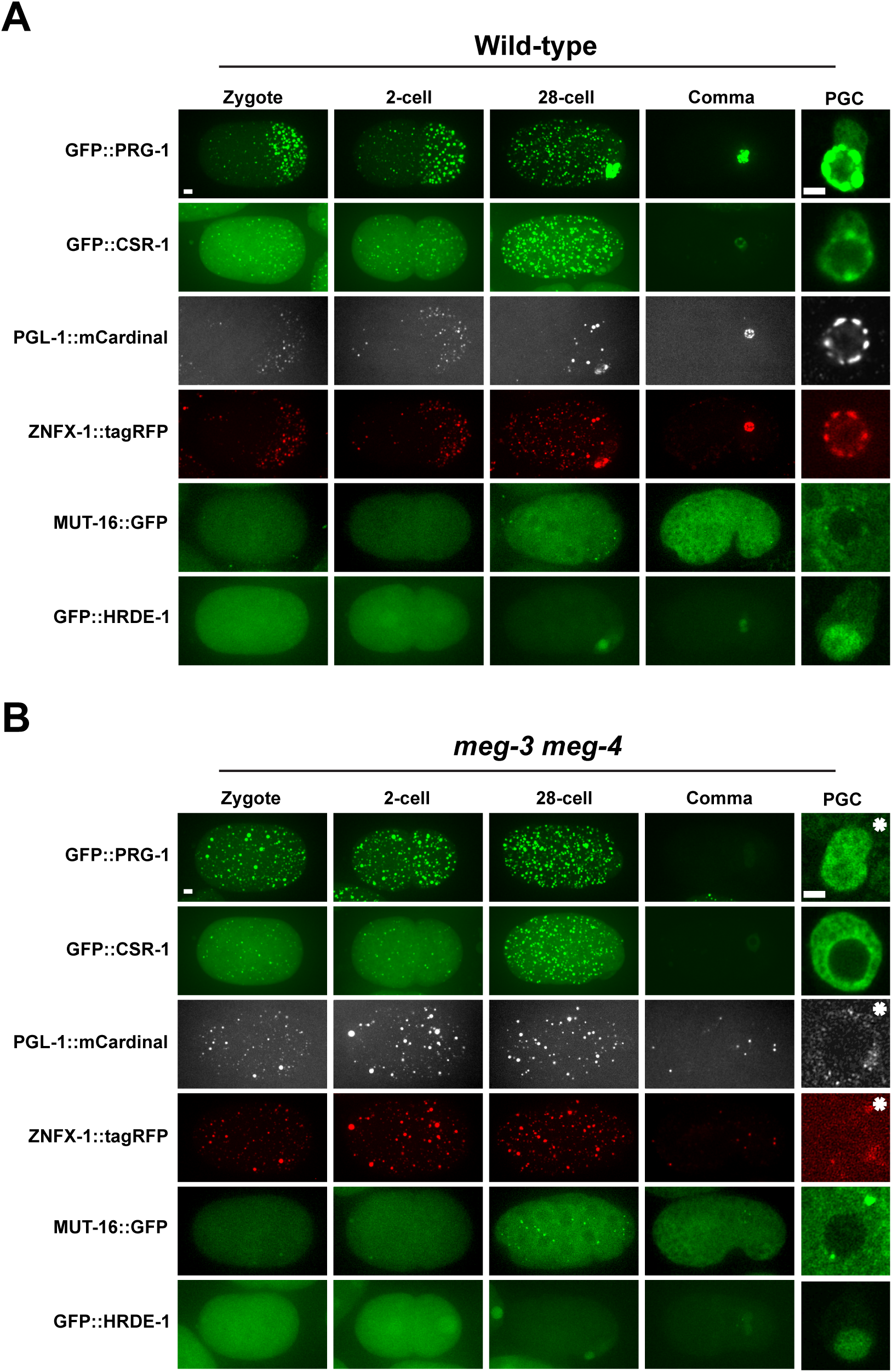
Localization of epigenetic factors during embryonic development in wild-type and *meg-3 meg-4* mutants. Photomicrographs of (A) wild-type and (B) *meg-3 meg-4* embryos at the indicated developmental stages expressing fluorescently-tagged nuage proteins and HRDE-1. All tags were introduced at the endogenous locus by genome editing. Last column shows close-ups of a single primordial germ cell (PGC) at comma-stage. Image acquisition and display values were adjusted for each protein and therefore levels cannot be compared between proteins. Wild-type and *meg-3 meg-4* panels for each fusion are comparable, except for panels with asterisks which were adjusted to visualize the much lower levels of fluorescence in *meg-3 meg-4* mutants. See Fig. S4F for non-adjusted panels. Scale bars are 4 µm (embryo panels) and 2 µm (PGC panels).

In *meg-3 meg-4* embryos, PGL-1, PRG-1, CSR-1 and ZNFX-1 granules were segregated evenly to all cells and turned over in somatic cells after the 28-cell stage (Fig. 5B). Consistent with failed preferential segregation to the germ lineage, by mid-embryogenesis (comma-stage), PGL-1, PRG-1, and ZNFX-1 levels were severely reduced in *meg-3 meg-4* compared to wild-type (Fig. 5B). In contrast, CSR-1 levels appear comparable to wild-type. At this stage, in wild-type, PGL-1, PRG-1, ZNFX-1 and CSR-1 are concentrated in granules around the nuclei of Z2 and Z3 (Fig. 5A). In contrast, in *meg-3 meg-4* mutants, these proteins were mostly cytoplasmic in Z2 and Z3 forming only rare puncta, with the exception of PGL-1 which formed many small cytoplasmic puncta (Fig. 5B).

Unlike P granule-associated proteins, the mutator foci protein MUT-16 was segregated uniformly to all cells of early wild-type embryos, and remained as an abundant cytoplasmic protein in most cells throughout embryogenesis. Bright perinuclear MUT-16 puncta could be observed in many cells, including Z2 and Z3. This pattern was not disrupted significantly in *meg-3 meg-4* mutants (Fig. 5B).

Finally, we also examined the embryonic distribution of HRDE-1, using a GFP-tagged allele (Methods). HRDE-1 was present in all cells in early embryos and became restricted to the germline founder cell P_4_ by the 28-cell stage by an unknown mechanism. This pattern was not disrupted in *meg-3 meg-4* embryos. In comma-stage embryos, HRDE-1 was present exclusively in Z2 and Z3 in both wild-type and *meg-3 meg-4* mutants (Fig. 5A-B). The only observed difference was that the nuclear-to-cytoplasm ratio of HRDE-1 was higher in *meg-3 meg-4* primordial germ cells compared to wild-type (Fig. 5A-B and S4G-H). No such difference was seen when comparing HRDE-1 in oocytes of *meg-3 meg-4* and wild-type hermaphrodites (Fig. S4G-H). Intriguingly, increased nuclear-to-cytoplasm ratio has been correlated with 22G-RNA loading for the somatic nuclear Argonaute, NRDE-3 (Guang et al., 2008).

In summary, we find that primordial germ cells in *meg-3 meg-4* mutants maintain mutator foci and nuclear HRDE-1, but fail to assemble perinuclear P and Z granules. P (PRG-1, CSR-1, PGL-1) and Z (ZNFX-1) granule proteins are still present in these cells, but are dispersed throughout the cytoplasm.

### *rde-11* and *sid-1* transcripts are transcribed and accumulate in P granules in wild-type, but not *meg-3 meg-4* primordial germ cells

The dramatic nuage assembly defect in *meg-3 meg-4* embryos led us to investigate whether *rde-11* and *sid-1* might be expressed in Z2 and Z3 during embryogenesis. We performed fluorescent *in situ* hybridization for *rde-11* and *sid-1* on wild-type embryos expressing GFP::PRG-1. Consistent with expression in the adult maternal germline, we detected cytoplasmic *rde-11* and *sid-1* transcripts in early embryos (Fig. S5). In comma-stage embryos, we observed scattered single *sid-1* and *rde-11* transcripts in somatic cells and clusters of *rde-11* and *sid-1* transcripts in Z2 and Z3 (Fig. S5). The clusters overlapped with perinuclear granules positive for GFP::PRG-1 (Fig. 6A). We also detected a few transcripts in the cytoplasm away from GFP::PRG-1 granules, but these were a minority (Fig. 6B). Consistent with zygotic transcription at this stage, we detected nuclear signal in 9 of 14 comma-stage embryos examined for *rde-11* expression and 4 of 5 comma-stage embryos examined for *sid-1* expression. These observations suggest that *rde-11* and *sid-1* are transcribed in Z2 and Z3 during embryogenesis and accumulate in P granules with PRG-1.

**Fig. 6:**
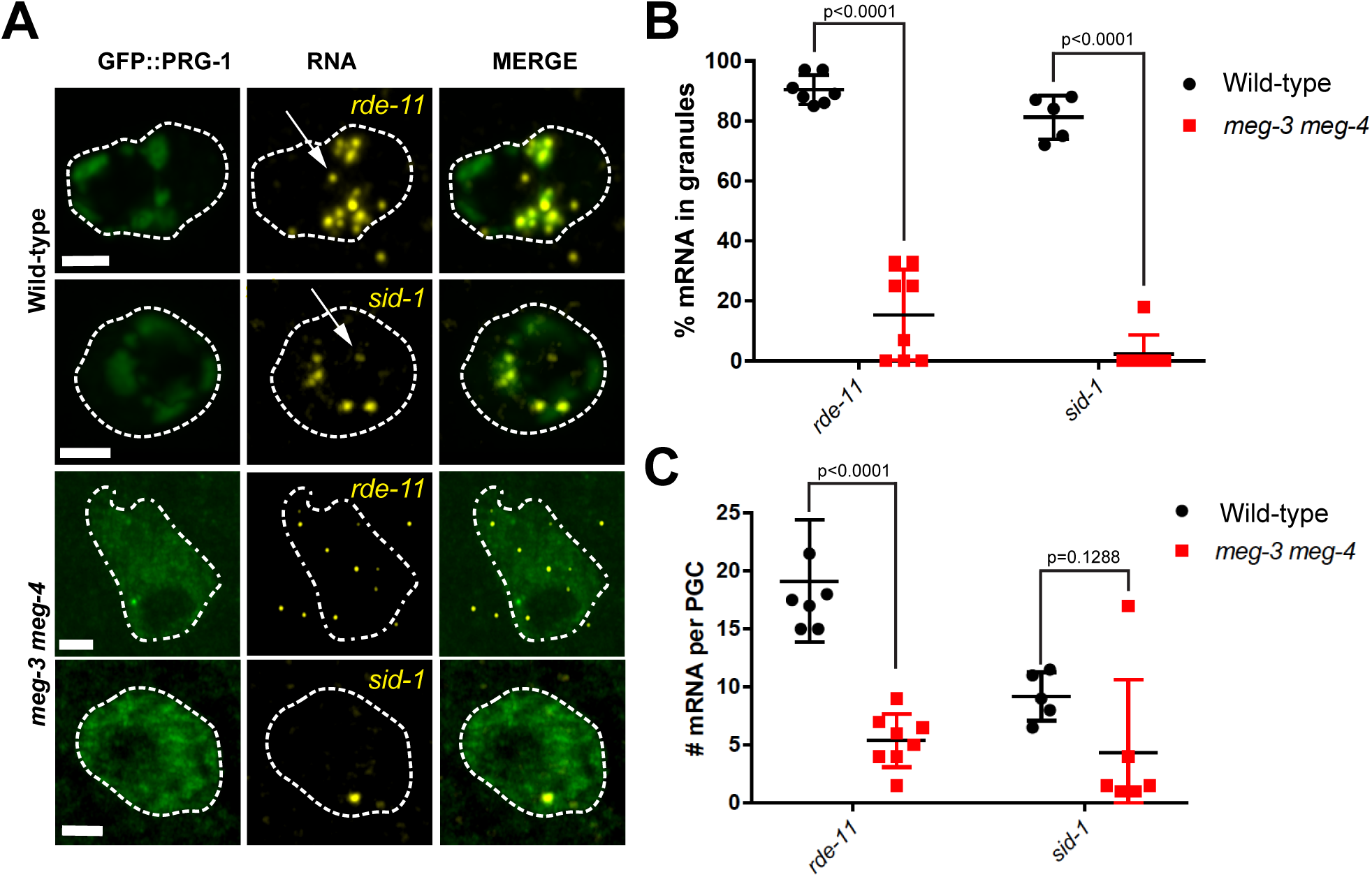
Localization of *rde-11* and *sid-1* transcripts in wild-type and *meg-3 meg-4* primordial germ cells. A. Photomicrographs of primordial germ cells in comma-stage embryos hybridized to fluorescent probes to visualize *rde-11* and *sid-1* transcripts (yellow). Embryos also express GFP::PRG-1 fusion (green). Arrows point to nuclear transcripts. Stippled lines indicate cell outline. Scale bar is 2 μm. B. Graph showing the % of *rde-11* and *sid-1* transcripts in GFP::PRG-1 granules in wild-type vs *meg-3 meg-4* primordial germ cells. Each dot represents one embryo. Error bars represent the standard deviation. P-values were obtained through an unpaired t-test. C. Graph showing the number of *rde-11* and *sid-1* transcripts in wild-type and *meg-3 meg-4* primordial germ cells. Each dot represents one embryo. Mid bar represents the mean while error bars indicate the standard deviation. P-values were obtained through an unpaired t-test. A significant p-value was obtained between mRNA number in wild-type and *meg-3 meg-4* for *rde-11* mRNA but was not for *sid-1* mRNA due to a single outlier.

Next, we examined *rde-11* and *sid-1* transcripts in *gfp::prg-1*; *meg-3 meg-4* embryos. *meg-3 meg-4* primordial germ cells accumulated fewer *rde-11* and *sid-1* transcripts compared to wild-type (Fig. 6A, C). We detected nuclear transcripts in 3 of 8 embryos examined for *rde-11* expression and 3 of 8 embryos examined for *sid-1* expression. Consistent with the fact that PRG-1 forms fewer and smaller granules in *meg-3 meg-4* mutants, a smaller proportion of cytoplasmic *rde-11* and *sid-1* transcripts were enriched in granules compared to wild-type and most transcripts were dispersed in the cytoplasm (Fig. 6A and B). We conclude that *rde-11* and *sid-1* loci are also transcribed in *meg-3 meg-4* primordial germ cells, albeit at a potentially lower efficiency compared to wild-type. *rde-11* and *sid-1* transcripts accumulate with PRG-1 in P granules in wild-type primordial germ cells, but not in *meg-3 meg-4* where they disperse with PRG-1 in the cytoplasm.

## DISCUSSION

In this study, we take advantage of a mutant deficient in nuage coalescence during embryogenesis to examine the function of nuage compartments in regulating endogenous gene expression. We find that *meg-3 meg-4* mutants become RNAi-deficient over several generations and that this phenotype requires PRG-1 and HRDE-1 activities. *meg-3 meg-4* mutants upregulate sRNAs against ∼300 loci, including four genes required for exogenous RNAi (*rde-11*, *sid-1, hda-3, zfp-1*) and two genes implicated in sRNA pathways (*wago-2* and *set-23*). The genes with upregulated sRNAs in *meg-3 meg-4* mutants belong to a unique class of loci that are targeted by PRG-1-piRNA and HRDE-1-sRNA complexes, and not targeted by CSR-1-sRNA complexes. *rde-11* and *sid-1* transcripts are expressed in primordial germ cells where they accumulate in perinuclear P granules in wild-type, but not in *meg-3 meg-4* mutants where the transcripts scatter in the cytoplasm mixing with other dispersed nuage components. Together, these observations suggest that coalescence of nuage into distinct condensates restrains 22G-RNA amplification initiated by piRNAs, especially at loci required for exogenous RNAi.

### Maternal inheritance of P granules is not essential for inheritance of epigenetic traits

In *Drosophila*, maternally-deposited piRNAs defend progeny against active transposable elements (Brennecke et al., 2008). Similarly, in *C. elegans*, maternal piRNAs are required to restore transposon silencing and the proper balance of 22G-RNAs in animals that do not inherit 22G-RNAs from their parents (Phillips et al., 2015; de Albuquerque 2015). How piRNAs and other sRNAs are transmitted from germline to germline across generations is not known. In principle, P granules (and their equivalent in *Drosophila*, the polar granules) are ideal conduits, since P granules concentrate Argonaute proteins and are actively partitioned to the embryonic germline during early embryonic cleavages. Our observations with *meg-3 meg-4* mutants, which break the cycle of maternal P granule inheritance, however, challenge this hypothesis. First, the fact that most germline genes are expressed normally in *meg-3 meg-4* mutants demonstrates that maternal inheritance of P granules is not essential to license most germline gene expression. Second, *meg-3 meg-4* become RNAi defective only after several generations, consistent with transmission of an epigenetic signal that is amplified over generational time. Finally, the RNAi-defective phenotype of *meg-3 meg-4* mutants is inherited maternally, providing direct evidence for epigenetic inheritance in the absence of embryonic P granules. We conclude that P granules are not essential to deliver epigenetic signals to the next generation. This conclusion does not exclude the possibility that some epigenetic signals may rely on embryonic P granules for maximal transmission (such as PRG-1/piRNAs complexes, see below).

The nuclear Argonaute HRDE-1 is likely to be the conduit for at least part of the epigenetic inheritance we observe in *meg-3 meg-4* mutants. HRDE-1 is required for the RNA-interference defect of *meg-3 meg-4* mutants. Nuclear HRDE-1 segregates with the embryonic germ line and this distribution was not affected in *meg-3 meg-4* mutants. CSR-1 and PRG-1 could also be detected in the cytoplasm of *meg-3 meg-4* primordial germ cells, despite not being in perinuclear condensates. These observations suggest that at least some of the maternal pool of Argonautes present in zygotes segregates with the embryonic germ lineage independent of P granules. In zygotes, the polarity regulators PAR-1 and MEX-5 collaborate to drive asymmetric segregation of germ plasm (a collection of maternally-inherited RNA-binding proteins) to the germline founder cell P_4_ (Schubert et al., 2000; Folkmann and Seydoux, 2019). It will be important to investigate the mechanisms that segregate HRDE-1 and other Argonautes to the embryonic germline and ensure transmission of epigenetic signals from one generation to the next.

### P granules protect *rde-11* and *sid-1* from PRG-1/HRDE-1-driven silencing

Several lines of evidence suggest that the RNAi deficient phenotype of *meg-3 meg-4* is due to silencing of genes required for exogenous RNAi, in particular *rde-11* and *sid-1*. First, like *rde-11* mutants (Zhang et al., 2012), *meg-3 meg-4* mutants exhibit both reduced production of secondary sRNAs in response to an exogenous trigger and reduced levels of endogenous sRNAs at 108 loci also affected in *rde-11* mutants. Second, like *sid-1* mutants (Wang and Hunter, 2017), *meg-3 meg-4* mutants are partially resistant not only to dsRNA introduced by feeding but also to dsRNA introduced by injection. Third, sRNAs mapping to the *rde-11* and *sid-1* loci were elevated in four independent *meg-3 meg-4* lines, and both transcripts were reduced in the original *meg-3 meg-4* line. Fourth, loss of *hrde-1* in *meg-3 meg-4* restored both competence for RNAi and *rde-11* transcript levels in adult gonads. Although silencing of *rde-11* and *sid-1* are likely to be the main drivers of the *meg-3 meg-4* RNAi-defective phenotype, they may not be the only contributors. sRNAs against two other genes required for RNAi (*hda-3* and *zfp-1*) and two genes implicated in sRNA pathways (*wago-2* and *set-23*) were also elevated in *meg-3 meg-4* strains. To what extent silencing of these and other genes additionally contributes to the *meg-3 meg-4* RNAi-defective phenotype remains to be determined.

Of the thousands of genes expressed in germ cells, what makes *rde-11* and *sid-1* so prone to silencing in *meg-3 meg-4* mutants? Examination of recent transcriptome-wide data for PRG-1/piRNA engagement on endogenous transcripts revealed that *rde-11* and *sid-1* are among the top 50 most targeted messages in the entire *C. elegans* transcriptome (Shen et al., 2018). In contrast, *rde-11* and *sid-1* do not appear to be targeted by sRNAs associated with the protective Argonaute CSR-1. This combination of excessive targeting by PRG-1 and hypo-targeting by CSR-1 may be a contributing factor for why *rde-11* and *sid-1* are selectively silenced in *meg-3 meg-4* mutants.

Another characteristic of *rde-11* and *sid-1* is that they are expressed in primordial germ cells during embryogenesis. Only three other genes so far have been documented to be transcribed in primordial germ cells before hatching (Subramaniam and Seydoux, 1999; Kawasaki et al., 1998; Mainpal et al., 2015), which has been described as a period of low transcriptional activity for the germline (Schaner et al., 2003). This is also precisely the developmental period during which *meg-3 meg-4* mutants lack P granules, suggesting that expression in the absence of P granules is what triggers silencing of *rde-11* and *sid-1* in *meg-3 meg-4* mutants.

We propose the following model (Fig. 7). In wild-type, upon emergence from the nucleus, *rde-11* and *sid-1* transcripts accumulate in P granules where they associate with PRG-1/piRNA complexes. Transcript retention in P granules limits their use as templates for 22G-RNA synthesis in Z granules and mutator foci. Consequently, only a moderate number of HRDE-1-associated 22Gs accumulate against *rde-11* and *sid-1* in wild-type, allowing the loci to remain expressed. In contrast, in *meg-3 meg-4* mutants, *rde-11* and *sid-1* transcripts are released directly in the cytoplasm where they are free to mix with dispersed nuage components. 22G-RNA synthesis is accelerated, causing HRDE-1 to become hyperloaded with sRNAs against *rde-11* and *sid-1,* enter the nucleus and silence the rde*-11* and *sid-1* loci. The observed increase in HRDE-1 nuclear-to-cytoplasmic ratio in *meg-3 meg-*4 primordial germ cells is suggestive of elevated HRDE-1 nuclear activity.

**Fig. 7:**
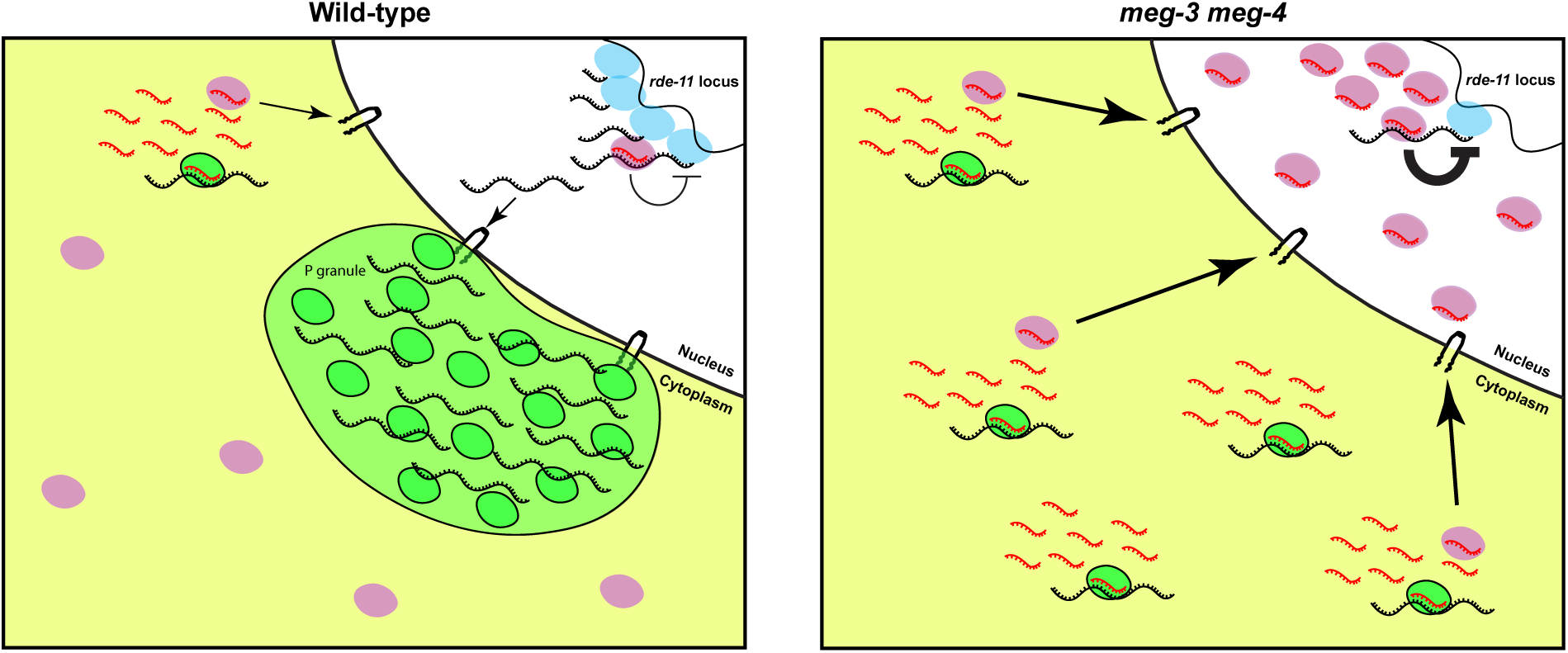
Model illustrating the fate of *rde-11* transcripts in wild-type and *meg-3 meg-4* primordial germ cells. In wild-type primordial germ cells, rde*-11* transcripts (black) are transcribed by RNA polymerase II (blue), and accumulate in P granules (green) upon exit from the nucleus. In P granules, *rde-11* transcripts are targeted by PRG-1/piRNA complexes (green) which slows their release into the cytoplasm. Few transcripts reach the cytoplasm (yellow) where mutator activity triggers production of secondary sRNAs (red) that load on HRDE-1 (pink). In *meg-3 meg-4* primordial germ cells, *rde-11* transcripts immediately disperse in the cytoplasm upon exit from the nucleus. In the cytoplasm, *rde-11* transcripts are targeted by PRG-1/piRNA complexes and by mutator activity which triggers the production of secondary sRNAs. The secondary sRNAs are loaded on HRDE-1 stimulating its nuclear accumulation leading to silencing of the *rde-11* locus.

It may appear counterintuitive that *rde-11* and *sid-1* transcripts experience an increase in PRG-1-driven silencing, given that PRG-1 levels are much lower overall in *meg-3 meg-4* primordial germ cells compared to wild-type (Fig. 5). In certain genetic contexts, maternal inheritance of PRG-1 has been shown to *protect* germline mRNAs from silencing by preventing misrouting of 22G-RNAs into silencing Argonaute complexes (Phillips et al., 2015). One possibility is that targeting by PRG-1/piRNA complexes in the context of the P granule environment marks transcripts for potential silencing but also protects them from mutator activity in the cytoplasm by retaining most transcripts in granules. In the absence of P granules, however, the protective influence of PRG-1/piRNA complexes is lost and transcripts are free to engage with the sRNA amplification machinery in the cytoplasm. The low levels of PRG-1 in *meg-3 meg-4* primordial germ cells may explain why several rounds of cytoplasmic exposure (generations) are needed before sufficiently high numbers of HRDE-1/sRNA complexes are generated to silence the RNAi genes.

### A mechanism for fine tuning the RNA-interference machinery?

piRNAs are genomically-encoded so presumably the heavy targeting of *rde-11* and *sid-1* is beneficial to *C. elegans*. The ability to mount an RNAi response in *C. elegans* has been reported to be tunable across generations (Houri-Ze’evi et al., 2016). Transgenerational duration of an RNAi response to a primary dsRNA trigger is extended when progeny are exposed to an unrelated second dsRNA trigger. Furthermore, exposure to dsRNA changes the level of sRNAs that target genes in the RNA-interference machinery, including *rde-11* and *sid-1* and many others (Houri-Ze’evi et al., 2016). Small changes in temperature have also been shown to affect piRNA biogenesis leading to changes in gene expression in subsequent generations (Belicard et al., 2018). These observations suggest that environmental influences can modulate the potency and specificity of the sRNA machinery. We suggest that this modulation is achieved in part by piRNA-targeting and sequestration in P granules of transcripts coding for epigenetic factors, such as *rde-11* and *sid-1*. An exciting possibility is that P granules modulate the rate of delivery of piRNA-targeted transcripts to mutator foci as a function of maternal experience and this process begins as soon as transcription initiates in the primordial germ cells. In this way, embryos could integrate ancestral inputs to fine-tune their own epigenetic machinery before hatching and taking their first meal.

## Acknowledgments

We thank the Johns Hopkins Neuroscience Research Multiphoton Imaging Core (NS050274) and the Johns Hopkins Integrated Imaging Center (S10OD023548) for excellent microscopy support. We thank the Phillips and Mello labs for strains, the Mello lab for help with analyzing CLASH data, the Miska lab for guidance with sRNAseq, Anne Dodson and Scott Kennedy for sharing data before publication, and Charlotte Choi, Anne Dodson, Scott Kennedy, John Kim, Craig Mello, Eric Miska, the Baltimore Worm Club and the Seydoux lab for many helpful discussions. JPTO thanks Taylor Swift for inspiration. JPTO was supported by the JHU SOM Biochemistry, Cellular, and Molecular Biology NIH training grant (T32 GM007445). GS is an investigator of the Howard Hughes Institute.

## Author Contributions

JPTO, AWF, LB conducted the experiments; JPTO, AWF, LB, CYL and GS analyzed the data; US, AGC, and JMC constructed the GFP::CSR-1 and GFP::HRDE-1 strains, JPTO and GS designed the experiments and wrote the paper.

## STAR Methods

### Lead Contact and Materials Availability

Further information and requests for resources and reagents should be directed to Geraldine Seydoux. Plasmids generated in this study have been deposited to Addgene. Strains used in this study have been deposited at the Caenorhabditis Genetics Center (CGC). Unique reagents generated in this study are listed in the Key Resources Table.

### Experimental Model and Subject Details

All *C. elegans* strains used throughout this study were maintained at 20° C on NNGM growth media or Enriched Peptone media and fed OP50 or NA22 bacteria. Strains used in this study are listed in the Key Resources Table.

### Methods Details

#### Strain construction and validation

CRISPR generated lines were created as in Paix et al., 2017 or Dickinson et al 2015 as indicated in the Key Resources Table. Guides and repair temples used for CRISPR are listed in Table S5. For functional validation of the *gfp::hrde-1* and *gfp::csr-1* strains, brood sizes were determined as follows: L4 stage worms were picked to separate plates and transferred every day until egg laying ceased. The progeny on each plate were counted 1-2 days after the mother was transferred. Experiments were conducted at 25° C and 20° C for *gfp::hrde-1* and *gfp::csr-1* respectively (Fig. S4D-E).

The following names were used throughout the paper to indicate the following strains:

- *meg-3 meg-4^#1^* → JH3475
- *meg-3 meg-4 ^#2^* → JH3672
- *meg-3 meg-4 ^#3^* → JH3673
- *meg-3 meg-4 ^#4^* → JH3674

#### RNA interference assays

The *pos-1* 400 nt L4440 RNAi vector used for sRNA sequencing in Fig 2C, D was made using the Clontech In-Fusion HD Cloning Kit. The PCR oligos used for cloning are listed in Table S5. The *pos-1* segment cloned was amplified from the full CDS *pos-1* L4440 plasmid from the Dharmacon *C. elegans* RNAi collection and cloned into the L4440 vector.

All RNAi experiments were performed at 20°C. Feeding RNAi experiments were performed by placing worms on HT115 bacteria expressing dsRNA as previously described in Timmons and Fire, 1998. Briefly, HT115 cells were transformed with L4440 RNAi plasmids, and colonies were inoculated into 2 mLs 100 ug/mL ampicillin LB liquid media and grown for five hours at 37° C. Cultures were then induced with IPTG for a final concentration of 5 mM and grown for 45 minutes. Bacteria were then plated on NNGM agar containing 100 ug/mL carbenicillin and 1 mM IPTG. Feeding was performed starting at the L1 or L4 stage (time of feeding is indicated in the figure legends for each experiment). For feeding at the L1 stage, worms were fed RNAi bacteria for ∼72 hours before experimentation. For feeding at the L4 stage, experiments were performed ∼36 hours after placement on RNAi.

For RNAi by injection, *pos-1* dsRNA was obtained using the T7 RiboMAX Express Large Scale RNA Production System and purified using Zymo’s RNA Clean & Concentrator Kits. Young adults were injected with 200 ng/uL *pos-1* dsRNA and embryonic lethality was assessed for each injected mother 16 hours following injection.

For embryonic lethality calculations, single mothers or cohorts of 10-20 mothers were allowed to lay eggs for periods ranging from 1-2 hours. Embryos were then counted, and adults were scored four days later*. prg-1; meg-3 meg-4* hermaphrodites lay ∼50% dead embryos even under non-RNAi conditions. For those experiments, embryonic lethality on *pos-1* RNAi was normalized to embryonic lethality on control L4440 RNAi.

#### Western Blots

For the MEG-3::OLLAS/MEG-4::3X::FLAG western blot, a mixed population of worms was subjected to bleaching to obtain embryos for L1 synchronization by shaking in M9 (22.0 mM KH_2_PO_4_, 42.3 mM Na_2_HPO_4_, 85.6 mM NaCl, 1 mM MgSO_4_) for 18-20 hours. L1 samples were then taken before plating on OP50 bacteria. Samples were then collected at different developmental stages. Embryo samples were collected from the synchronized gravid adult worms. Staged samples were resuspended in 1x PBS/cOmmplete Mini, EDTA-free Protease Inhibitor Cocktail. 5.5 uL of dense worm volume was then combined with 2.5 uL of NuPAGE LDS Sample Buffer and 2 uL of 1 M DTT.

For GFP::PRG-1/GFP::CSR-1 western blots, 75-100 fertile adults were collected and placed in 20 uL of 1x PBS/ cOmmplete Mini, EDTA-free Protease Inhibitor Cocktail. 9.09 uL of NuPAGE LDS Sample Buffer and 7.27 of 1 M DTT were added to each sample.

For sample preparation, all samples were lysed by four freeze thaw cycles. Following lysis, samples were heated at 85 C° for 10 minutes and then run on a Bolt 4-12% Bis-Tris Plus Gel in NuPAGE MOPS SDS Running Buffer. Samples were then transferred to an Immobilon-P PVDF Membrane, blocked in PBS+0.1%Tween20+5% nonfat dry milk and incubated with primary antibodies diluted in PBS+0.1%Tween20+5% milk. The blot was washed three times in PBS+0.1%Tween20 and visualized by treatment with HyGLO Quick Spray Chemiluminescent HRP Antibody Detection Reagent and imaging by the KwikQuantTM Imager. For samples requiring a secondary antibody, the blot was incubated with a secondary antibody diluted in PBS+0.1%Tween20+5% milk following the three washes after the primary antibody. The blot was washed thrice more in PBS+0.1%Tween20 and imaged as described above.

Antibody dilutions used were as follows:

- anti-FLAG M2 mouse IgG1: 1:500 dilution
- anti-OLLAS L2 rat IgG1 Kappa HRP: 1:1000
- anti-α-Tubulin mouse IgG1: 1:1000
- anti-GFP mouse IgG2a: 1:500
- goat anti-mouse IgG1 HRP: 1:2500
- goat anti-mouse IgG2a HRP: 2500

#### RNA extraction and high-throughput sequencing library preparation

Mixed or adult staged (∼55-60 hours following L1 synchronization) populations of worms were collected, and RNA was isolated using the TRIzol reagent and chloroform. RNA was then concentrated and purified using Zymo’s RNA Clean & Concentrator Kits. For sRNA library preparation, RNA was either treated or untreated with RNA 5’ polyphosphatse (20 U/ug of RNA). Samples were then incubated for 30 minutes at 37° C and purified via phenol/chloroform extraction and ethanol precipitation supplemented with sodium acetate and glycogen. sRNA libraries were then constructed using 1 ug of polyphosphatase-treated/untreated total RNA as input into the TruSeq Small RNA Library Preparation Kit with 11 cycles of PCR amplification. Libraries were then size selected on a Novex 6% TBE gel and purified.

For mRNA sequencing, 1 ug of total RNA was treated with Ribo-Zero Gold Epidemiology rRNA Removal Kit. A 1:100 dilution of ERCC RNA Spike-In Mix was added. Libraries were then prepared using the TruSeq RNA Library Prep Kit v2 with 13 cycles of PCR amplification.

All sequencing was performed using the Illumina HiSeq2500 at the Johns Hopkins University School of Medicine Genetic Resources Core Facility.

#### High-throughput sequencing analyses

*sRNA sequencing*: 5’ sequencing adapters were trimmed using Cutadapt with default settings (Martin, 2011). Reads longer than 30 nts and shorter than 18 nts were discarded. Reads were then aligned to the UCSC ce10 *C. elegans* reference genome using Bowtie 2 (Langmead and Salzberg, 2012). Reads mapping to genetic features were counted using HTSeq-count (Anders et al., 2015) and differential expression analysis was conducted using DESeq2 (Love et al., 2014). For all our sRNA analysis, reads were normalized based on library size.

For sRNA class analyses, piRNA and miRNA lists were downloaded from WormBase. All other sRNAs were placed in Argonaute classes based on the locus targeted and published lists of loci targeted by sRNAs immunoprecipitated with specific Argonautes from wild-type worm lysates [Gu et al., 2009 (WAGO-1 IP), Xu et al, 2018 (WAGO-4 IP), Buckley et al., 2012 (HRDE-1 IP), and Claycomb et al., 2009 (CSR-1 IP)].

*mRNA sequencing* sequencing reads were aligned to the UCSC ce10 *C. elegans* reference genome usingHISAT2 (Kim et al., 2015). Reads aligning to genetic features were then counted using HTSeq-count (Anders et al., 2015) and analyzed for differential expression analysis using DESeq2 (Love et al., 2014).

Genome browser views were adapted from IGV TDF file visualization with zoom levels set to 7, window function set to “Mean,” and window size set to 5 (Robinson et al., 2011).

A list of high-throughput sequencing libraries generated in this study is listed in Table S6.

#### Single molecule fluorescence in situ hybridization (smFISH)

smFISH probes for *rde-11* and *sid-1* were designed using Biosearch Technologies’s Stellaris Probe Designer. The fluorophores used in this study were Quasar570 and Quasar670. For sample preparation, embryos or adult germlines were extruded from adults on poly-lysine slides and subjected to freeze-crack followed by methanol fixation. Samples were washed five times in PBS+0.1%Tween20 and fixed in 4% PFA for one hour at room temperature. Samples were again washed in PBS+0.1%Tween20 four times, twice in 2x SCC, and once in wash buffer (10% formamide, 2x SCC) before blocking in hybridization buffer (10% formamide, 2x SCC, 200 ug/mL BSA, 2mM Ribonucleoside Vanadyl Complex, 0.2 mg/mL yeast total RNA, 10% dextran sulfate) for 30 minutes at 37° C. Hybridization was then conducted by incubating samples with 50 nM probe solution diluted in hybridization buffer overnight at 37° C. Following hybridization, samples were washed twice in wash buffer at 37° C, twice in 2x SCC, once in PBS+0.1%Tween20, and twice in PBS. Lastly, samples were mounted using VECTASHIELD Antifade Mounting Media with DAPI or Prolong Diamond Antifade Mountant.

#### Microscopy

Fluorescence confocal microscopy was performed using a Zeiss Axio Imager with a Yokogawa spinning-disc confocal scanner. Images were taken using Slidebook v6.0 software (Intelligent Imaging Innovations) using a 63x objective. For imaging of primordial germ cells, fluorescence super-resolution microscopy was performed using ZEISS LSM 880-AiryScan (Carl Zeiss) equipped with a 63X objective. Images were acquired and processed using ZEN imaging software (Carl Zeiss). Equally normalized images were exported via either Slidebook v6.0 or ZEN, and contrasts of images were equally adjusted between control and experimental sets using ImageJ.

### Quantification and Statistical Analysis

Statistical analysis used in Figs 2, 4, S4, and 6 were performed using an unpaired t-test. Statistics for differential expression analysis were done using DESeq2 (Love et al., 2014).

FIJI was used for western blot quantification, *rde-11* smFISH signal quantification in the germline, and quantification of GFP::HRDE-1’s nuclear to cytoplasmic ratio. For western blot quantification, ROIs of constant area were placed over the GFP and tubulin bands and the integrated density values were measured. The ratios between GFP signal and tubulin signal was then calculated. For the *rde-11* germline quantification, ROIs were drawn in the late pachytene region of the germline and mean intensity values were calculated using maximum projection images. Unstained germlines were then used for background calculation, which was then subtracted from the calculated mean intensity of the germlines with probes. These values were then normalized to the average of wild-type and plotted accordingly. For GFP::HRDE-1 nuclear to cytoplasmic ratio in the -2 oocyte, germlines were extruded and single plane images were taken of the -2 oocyte. ROIs were drawn in the nucleus and cytoplasm, and the ratio of the mean intensities was calculated for wild-type and *meg-3 meg-4*. For GFP::HRDE-1 nuclear to cytoplasmic ratio in the PGCs, single plane images were taken of wild-type and *meg-3 meg-4* embryos at comma-stage. In a similar manner to the adult germline, the mean intensities of the nucleus and cytoplasm were calculated and compared in a ratio. smFISH quantification of PGC granule enrichment was conducted using Imaris Image Analysis Software visualization in 3D space. RNAs were counted manually, and the percentage localized in a GFP::PRG-1 granule was calculated.

### Data and Code Availability

Sequencing data has been deposited onto the Gene Expression Omnibus (GEO) and can be found using the following accession numbers:

XXXXXXXXXXXXXX

XXXXXXXXXXXXXX

XXXXXXXXXXXXXX

The *prg-1* sRNA sequencing data from Fig. S3D was obtained from SRR513312 (Lee et al., 2012) and its corresponding wild-type from SRR6691711 (Tang et al., 2018).

## Supplemental Information

**Fig. S1: related to Fig. 2.**
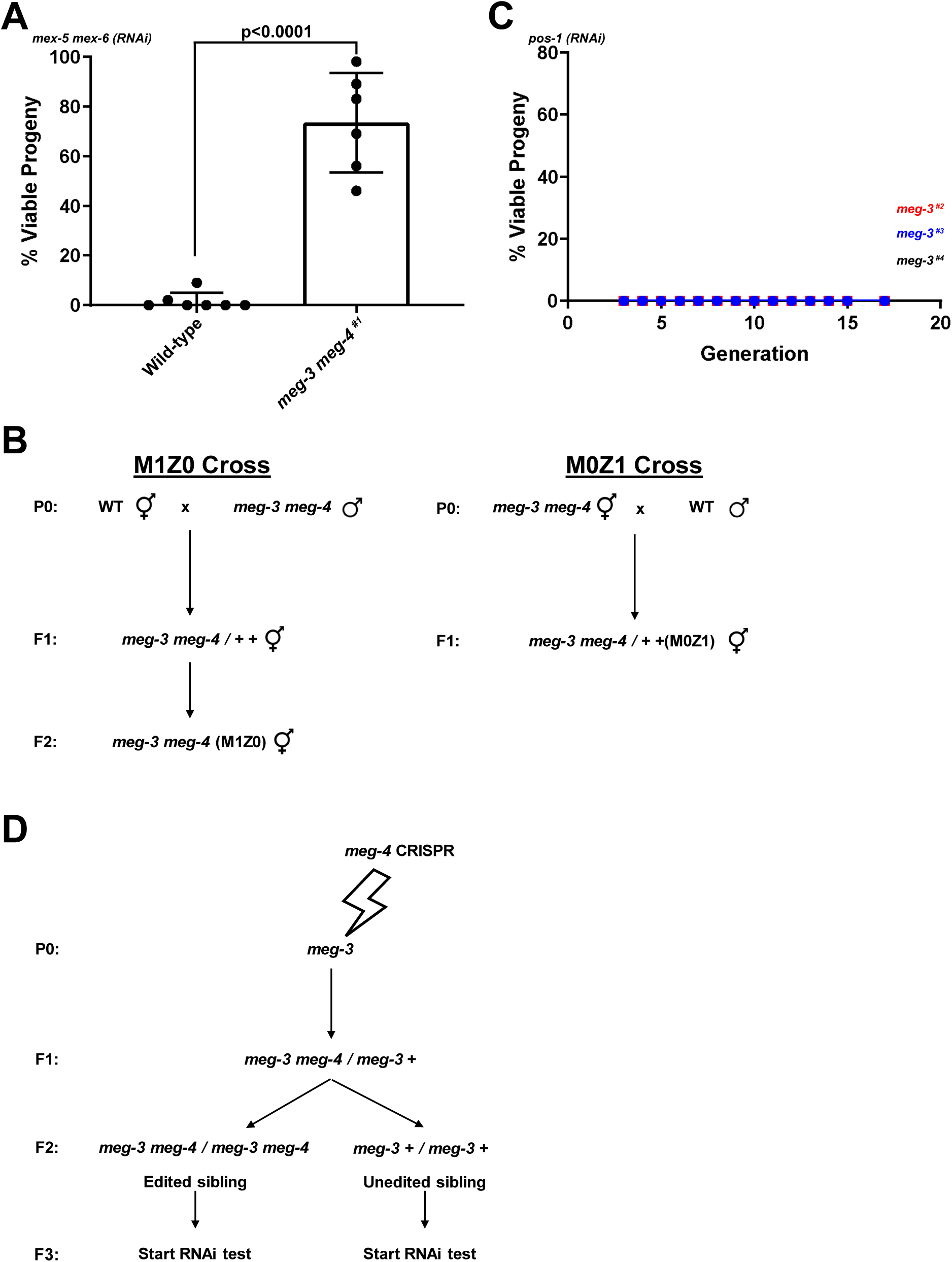
A. Graph showing the percentage of viable embryos among broods laid by ∼20 hermaphrodites of indicated genotypes fed with bacteria expressing *mex-5* and *mex-6* dsRNA from the L1 stage. Bar height represents the mean; error bars represent the standard deviation; the p-value was calculated using an unpaired t-test. B. Crosses used to generate hermaphrodites with varying numbers of maternal and zygotic *meg-3 meg-4* alleles. *meg-3 meg-4^#1^* hermaphrodites and males were used in all crosses. C. Graph showing the percentage of viable embryos among broods laid by ∼12 hermaphrodites carrying a deletion at the *meg-3* locus. The three strains shown were generated by cloning non-edited siblings of the *meg-3 meg-4* hermaphrodites analyzed in Fig. 2B. See S1D for CRISPR scheme. Unlike the *meg-3 meg-4* strains, all three *meg-3* strains exhibited complete RNAi penetrance (no viable progeny) throughout the experiment. D. Genome editing scheme to generate new *meg-3 meg-4* double deletion strains (and control sibling strains) from a strain carrying a deletion in *meg-3*. A single F1 animal was used to establish each *meg-3 meg-4* strain and its control sibling strain. F1 is Generation 1 in Figure 2B.

**Fig. S2: related to Fig. 3.**
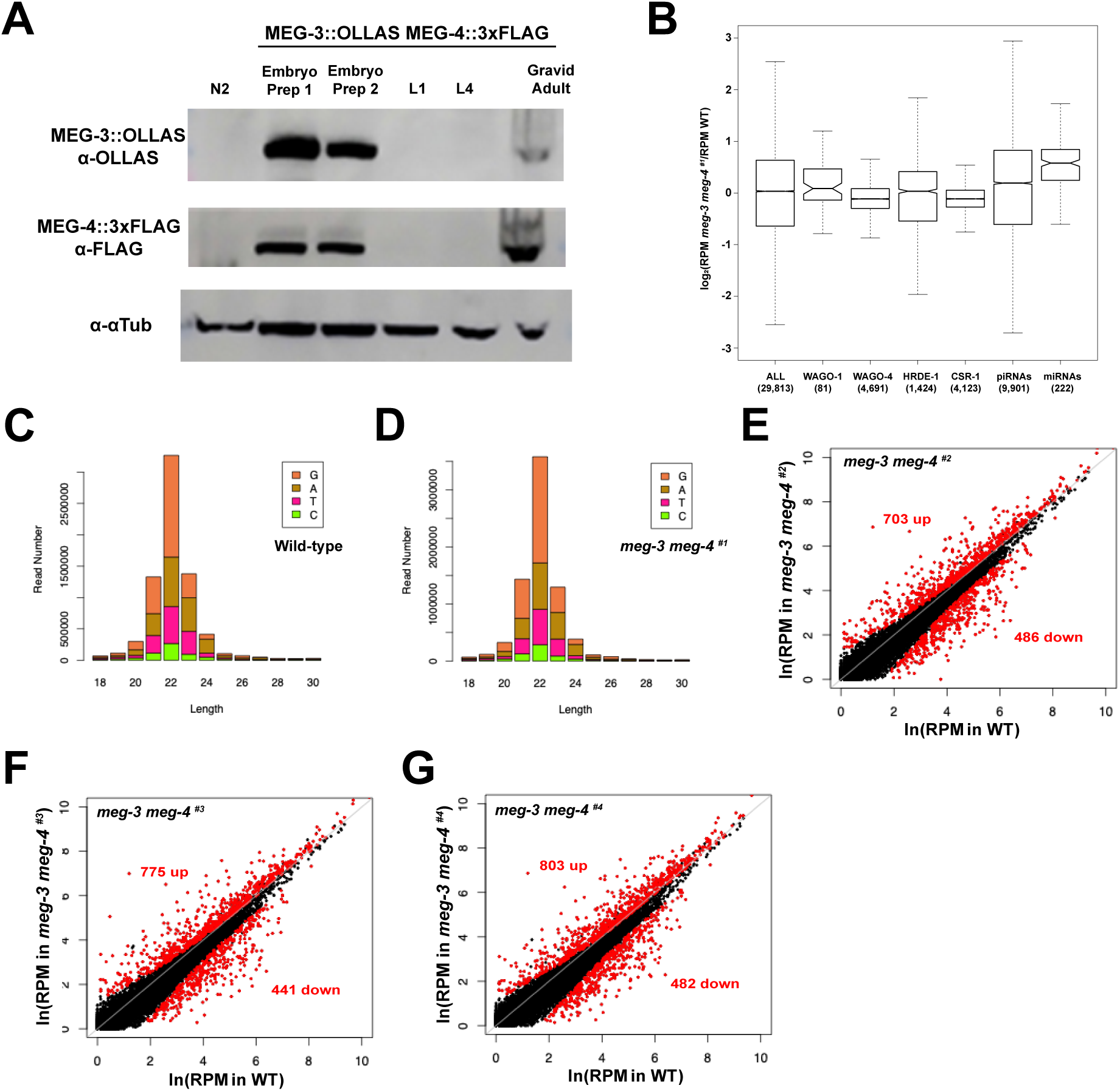
A. Western blot showing MEG-3 and MEG-4 protein levels in lysates collected at different developmental stages. Proteins were visualized using antibodies against epitope tags inserted at the *meg-3* and *meg-4* loci by genome editing. N2 refers to wild-type worms which do not contain epitope tags at the *meg-3* and *meg-4* loci. All other lanes were loaded with lysates prepared from worms in which *meg-3* and *meg-4* loci were tagged with OLLAS and 3xFLAG, respectively. Embryo Prep 1 lysate was prepared from embryos collected from 1-day old synchronized hermaphrodites. Embryo Prep 2 lysate was prepared from embryos collected from 1 to 3-day old hermaphrodites. L1 and L4 are first and fourth larval stages, respectively, and contain no embryos. Gravid adults contain embryos. Tubulin is used here as a loading control. B. Box plots showing the log2 fold change in abundance for the indicated classes of sRNAs in *meg-3 meg-4^#1^*animals compared to wild-type (Gu et al., 2009; Xu et al., 2018; Buckley et al., 2012; Claycomb et al., 2009; WormBase WS270). Boxes indicate the interquartile range; whiskers indicate the upper and lower quartiles; lines within the boxes indicate the median; notches display the confidence interval around the median. Parenthetical numbers indicate the number of sRNA-mapping genes represented by the respective box. Note that the WAGO-1 sRNA class as reported by Gu et al., 2009 only includes the ∼80 highest ranked sRNAs that immunoprecipitated with WAGO-1. As such, the WAGO-1 class of sRNAs may be underrepresented in our analysis. C-D. sRNA length distribution and 5’ nucleotide preference of sRNAs in wild-type and *meg-3 meg-4^#1^*. E-G. Scatter plots comparing sRNA abundance in wild-type (X-axis) and *meg-3 meg-4* (Y-axis) hermaphrodites in the indicated *meg-3 meg-4* strains. Each dot represents an annotated locus in the *C. elegans* genome. Red dots represent loci with significantly upregulated or downregulated sRNAs as determined from analysis using two biological replicates for each genotype.

**Fig. S3: related to Fig. 4.**
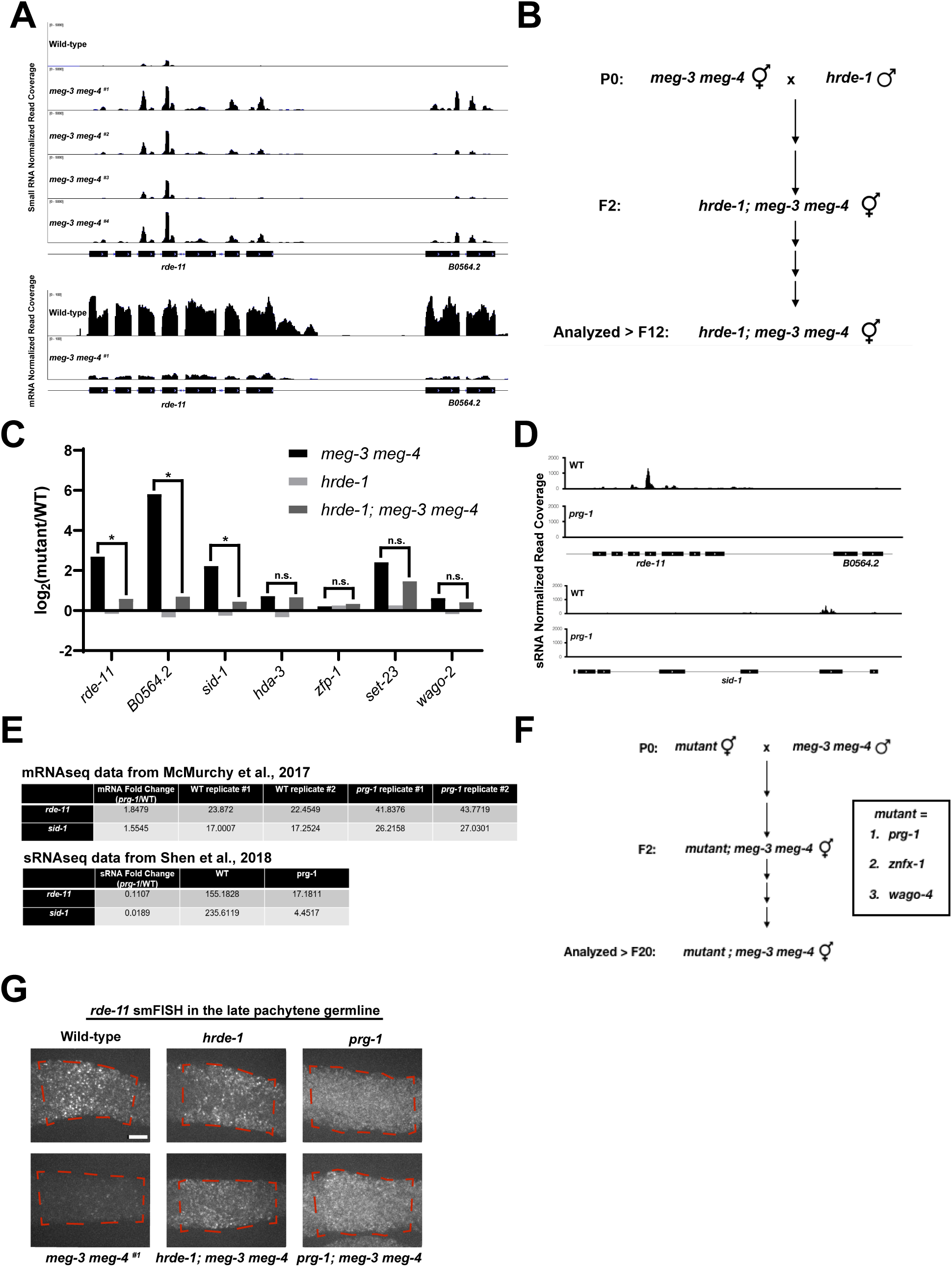
A. Browser view of the *rde-11/B0564.2* locus showing normalized sRNA reads in wild-type, *meg-3 meg-4^#1^*, *meg-3 meg-4 ^#2^*, *meg-3 meg-4 ^#3^*, and *meg-3 meg-4 ^#4^* mixed population. Lower panel shows mRNAseq data from *meg-3 meg-4^#1^* adults. Note the increase in sRNA reads and decrease in mRNA reads at both loci in *meg-3 meg-4^#1^* worms. B. Crosses used to generate *hrde-1; meg-3 meg-4* triple mutant. Genotypes were determined by PCR. C. Bar graph showing the log2 fold change in sRNAs mapping to the indicated loci in the indicated genotypes compared to wild-type. The graphs represent the log2 fold change from two biological replicates for each genotype. Stars indicate statistical significance in the comparison between *meg-3 meg-4* and *hrde-1; meg-3 meg-4* by DESeq2. D. Browser view of the *rde-11/B0564.2* and *sid-1* loci showing normalized sRNA reads in the indicated genotypes. Data from Tang et al., 2018/Lee et al., 2012. At both loci, sRNAs decrease in *prg-1* mutants. E. mRNA and sRNA fold changes and RPM values in *prg-1* compared to wild-type at the *sid-1*, *rde-11* loci from the indicated published studies. F. Crosses used to generate *prg-1; meg-3 meg-4*, *znfx-1; meg-3 meg-4*, and *wago-4; meg-3 meg-4 strains*. Genotypes were determined by PCR. G. Maximum projection photomicrographs of adult gonads of the indicated genotypes hybridized to fluorescent probes to visualize *rde-11* transcripts (quantified in Fig. 4C). Red stippled lines highlight the late pachytene region. Scale bar represents 10 μm.

**Fig. S4: related to Fig. 5.**
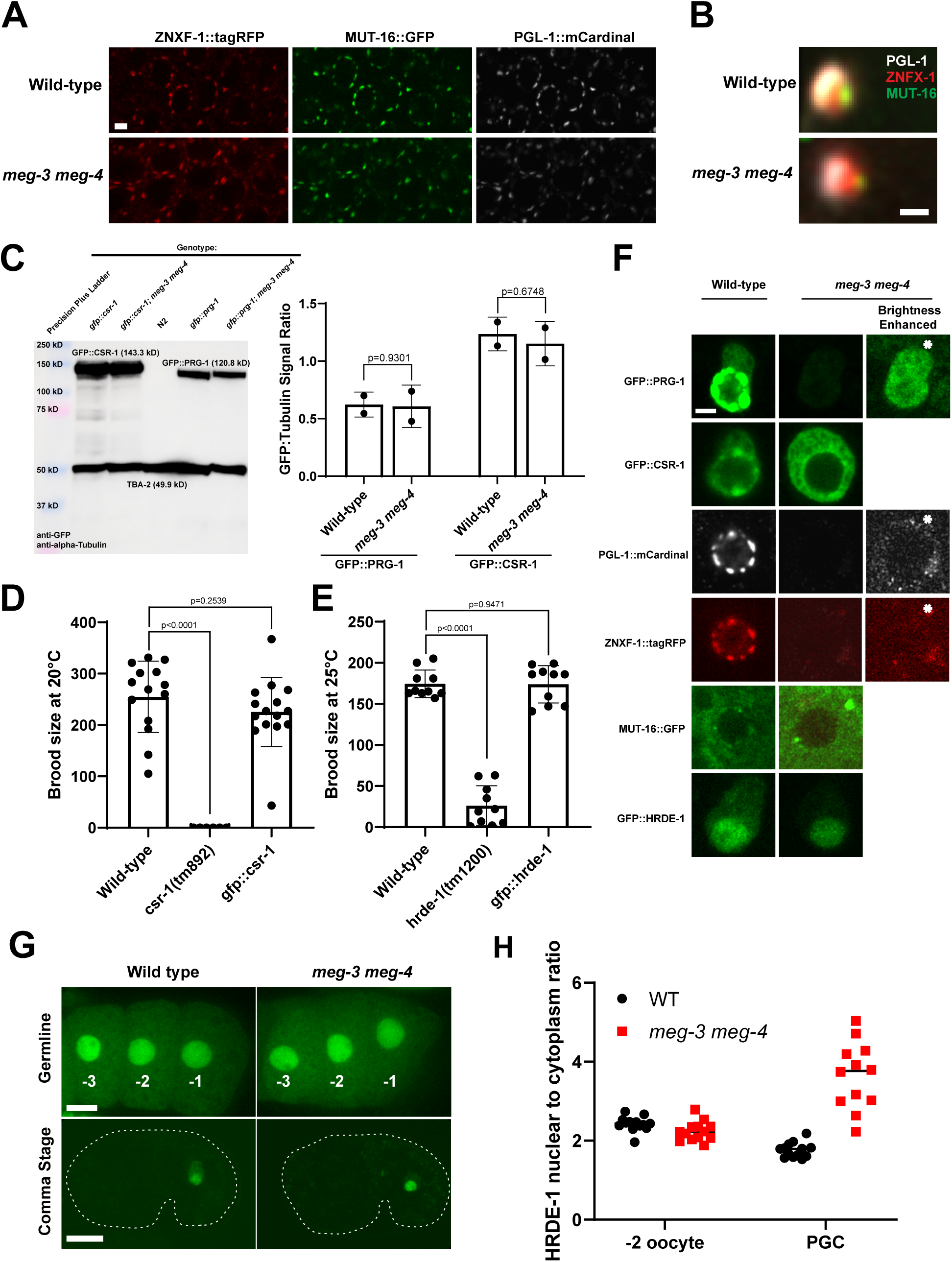
A. Photomicrographs showing germ cell nuclei (pachytene stage) in adult hermaphrodites of the indicated genotypes. No difference in the distribution in ZNFX-1, MUT-16 or PGL-1 are visible between wild-type and *meg-3 meg-4* at this stage. Scale bar is 2 µm. B. Same as A, but close-up showing the P granule - Z granule - Mutator pattern reported in Wan et al., 2018. Same pattern is visible in both wild-type and *meg-3 meg-4* strains. Scale bar is 500 nm. C. Anti-GFP/anti-α-Tub western blot of the indicated strains to assess levels of GFP::PRG-1 and GFP::CSR-1 in wild-type vs *meg-3 meg-4* adults. The ratio of GFP:Tubulin signal was measured and plotted accordingly. Bar height indicates the mean value; error bars represent the standard deviation; p-values were calculated using an unpaired t-test. Only fertile *meg-3 meg-4* adult worms were used in this analysis. No significance differences in PRG-1 and CSR-1 levels were detected. D-E. Functional validation of the *gfp::csr-1* and *gfp::hrde-1* lines. Strains were grown at 20° C and 25° C respectively and brood sizes were measured. Loss of function alleles for *csr-1* and *hrde-1* were included as a reference. Bar height indicates the mean value; error bars represent the standard deviation; p-values were calculated using an unpaired t-test. F. Photomicrographs (also shown in Fig. 5A-B) of single primordial germ cells at comma-stage to show unadjusted *meg-3 meg-4* panels (last row). Acquisition and display parameters for panels in first and second rows are identical, and demonstrate the lower levels of PRG-1/ZNFX-1/PGL-1 in *meg-3 meg-4* compared to wild-type. Panels in the last row (asterisk) have been enhanced for brightness to reveal the distribution of the low levels of PRG-1/ZNFX-1/PGL-1 in *meg-3 meg-4* mutants. Scale bar is 2 µm. G. Photomicrographs showing GFP::HRDE-1 in oocytes of adult hermaphrodites (top row) and in primordial germ cells of comma-stage embryos (bottom row) comparing wild-type and *meg-3 meg-4*. White stippled lines indicate cell outline. Scale bar represents 10 μm in both cases. H. Quantitation of the nuclear-to-cytoplasmic ratio of GFP::HRDE-1. Note the higher ratio in *meg-3 meg-4* primordial germ cells (PGC). Each dot represents one oocyte or one embryo.

**Fig. S5: related to Fig. 6.**
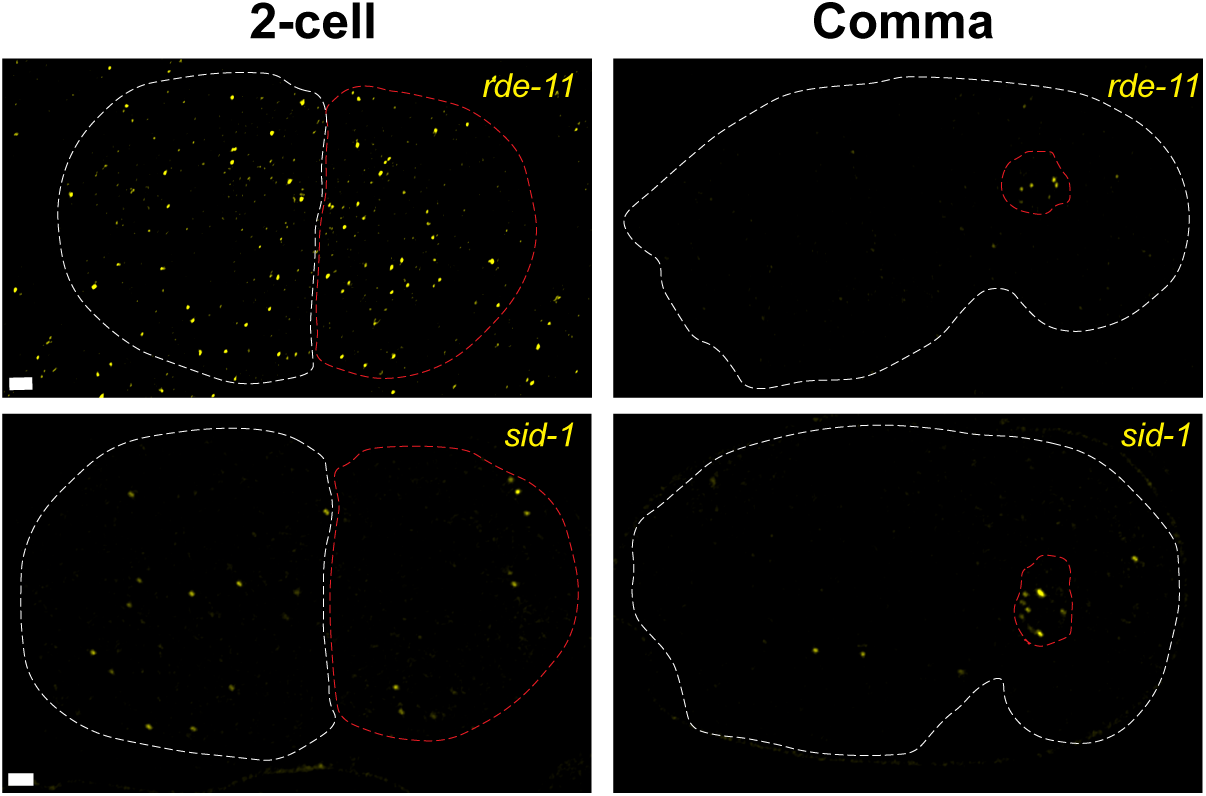
Photomicrographs of *pgl-1::gfp* embryos hybridized to fluorescent probes to visualize *rde-11* and *sid-1* transcripts (GFP not shown). Transcripts in 2-cell embryos represent maternal transcripts (white stippled cell is somatic blastomere AB, red stippled cell is germline blastomere P_1_). Transcripts in later comma-stage embryos are likely zygotic transcripts. Stippled white lines indicate embryo outline, stippled red lines indicate germ cell outline. Scale bar is 2 µm.

**Table S1.** Genes with sRNAs up in all four *meg-3 meg-4* strains.

**Table S2.** Genes with sRNAs down in all four *meg-3 meg-4* strains.

**Table S3.** Genes involved in RNAi.

**Table S4.** Misregulated *meg-3 meg-4* sRNAs rescued in *hrde-1; meg-3 meg-4*.

**Table S5.** List of guides, repair templates, and oligos used in this study.

**Table S6.** List of high-throughput sequencing libraries used in this study.

